# Interplay of EXO70 and MLO proteins modulates trichome cell wall composition and powdery mildew susceptibility

**DOI:** 10.1101/2022.12.30.521597

**Authors:** Jan W. Huebbers, George A. Caldarescu, Zdeňka Kubátová, Peter Sabol, Sophie C. J. Levecque, Hannah Kuhn, Ivan Kulich, Anja Reinstädler, Kim Büttgen, Alba Manga-Robles, Hugo Mélida, Markus Pauly, Ralph Panstruga, Viktor Žárský

## Abstract

EXO70 proteins are essential constituents of the octameric exocyst complex implicated in vesicle tethering during exocytosis, while MLO proteins are plant-specific calcium channels of which some isoforms play a key role during fungal powdery mildew pathogenesis. We here detected by a variety of histochemical staining procedures an unexpected phenotypic overlap of *A. thaliana exo70H4* and *mlo2 mlo6 mlo12* triple mutant plants regarding the biogenesis of leaf trichome secondary cell walls. Biochemical and Fourier transform infrared spectroscopic analyses of isolated trichomes corroborated deficiencies in the composition of trichome cell walls in *exo70H4* and *mlo2 mlo6 mlo12* mutants. Transgenic lines expressing fluorophore-tagged EXO70H4 and MLO variants exhibited extensive co-localization of these proteins at the trichome plasma membrane and cell wall. Furthermore, mCherry- EXO70H4 mislocalized in trichomes of the *mlo* triple mutant and, *vice versa*, MLO6- GFP exhibited aberrant subcellular localization in trichomes of the *exo70H4* mutant. Transgenic expression of GFP-marked PMR4 callose synthase, a previously identified cargo of EXO70H4 dependent exocytosis, revealed reduced cell wall delivery of GFP- PMR4 in *mlo* triple mutant plants. *In vivo* protein-protein interaction assays uncovered isoform-preferential physical interaction between EXO70 and MLO proteins. Finally, *exo70H4* and *mlo* mutants, when combined, showed synergistically enhanced resistance to powdery mildew attack. Taken together, our data point to an isoform-specific interplay of EXO70 and MLO proteins in the modulation of trichome cell wall biogenesis and powdery mildew susceptibility, possibly by (co-)regulating focal secretion of cell wall-related cargo.

## Introduction

Exocyst is an evolutionarily conserved multisubunit protein complex found in all eukaryotes. It consists of eight subunits (SEC3, SEC5, SEC6, SEC8, SEC10, SEC15, EXO70, EXO84) that together form a heterooligomeric complex (TerBush et al., 1996). Initially described in yeast (TerBush et al., 1996), it was later found in all eukaryotes including plants (Elias et al., 2003). Exocyst is a CATCHR type vesicle tethering complex that is especially involved in the tethering and docking of *post*-Golgi secretory vesicles to the plasma membrane (PM) and the regulation of soluble *N*- ethylmaleimide-sensitive-factor attachment receptor (SNARE) complex formation, which is decisive for membrane fusion events, including vesicle fusion at the PM (Heider and Munson, 2012). In addition to this conserved core function, individual subunits have been ascribed contributions to cellular processes such as autophagy and cytoskeleton regulation (Heider and Munson 2012).

In eukaryotes other than land plants, exocyst subunits are typically present as solitary isoforms encoded by single-copy genes. Differing from this scenario, the EXO70 subunit family expanded and diverged in land plants into three subfamilies with rapidly evolving paralogues (Synek et al., 2006; Cvrčková et al., 2012; Žárský et al., 2020). For example, in the dicotyledonous reference plant *Arabidopsis thaliana* there are 23 EXO70 isoforms and in the monocotyledonous reference crop plant rice more than 40 isoforms (Cvrčková et al., 2012). The EXO70 subunit is crucial for the membrane attachment of the entire exocyst complex due to its interaction with membrane lipids (Synek et al., 2021) and association with small GTPases of the Rho family (Rossi et al., 2020). Among other functions, plant EXO70 isoforms are involved in the localized secretion of cell wall components at the PM in root epidermis, seed coats, pollen tubes, and trichomes (Kubátová et al., 2019; Kulich et al., 2018; Sekereš et al., 2017; Fendrych et al., 2013; Kulich et al., 2010). Associated with this secretory pathway function, the exocyst complex, and especially specific EXO70 isoforms, participate in cellular defense/innate immunity. In this context, they contribute significantly to the biogenesis of defensive cell wall appositions (papillae) and the formation of perifungal membranes (Pecenková et al., 2017; Žárský et al., 2020). It is, therefore, not surprising that they are often targeted by pathogen effector proteins to overcome plant host immunity (Žárský et al., 2020; Pecenková et al., 2017; Fujisaki et al., 2015).

Trichomes are unbranched or branched polarized outgrowths of epidermal cells that are distributed on the surface of plant organs including leaves. In the case of the branched single-celled trichomes of *A. thaliana* rosette leaves, their development is genetically well-characterized (Hülskamp et al. 1994; Folkers et al. 1997). It can be divided into six major stages: initiation (including DNA endoreduplication), polar expansion, branching, branch growth, diffuse growth, and cellular maturation (Szymanski et al. 1998). *EXO70H4* (*At3g09520*) is the eleventh most highly expressed gene in mature trichomes (Jakoby et al., 2008). Recent studies (Kulich et al., 2015; Kulich et al., 2018) showed that EXO70H4 has an important role in trichome maturation, in particular in the deposition of secondary cell walls, especially in the apical region. Mature trichomes of *exo70H4* loss-of-function mutants appear to be more flexible than wild-type (WT) trichomes. This phenotype is due to the thinner secondary cell wall of mutant trichomes and because of the absence of the entire inner callose-rich cell wall layer, leading to a decrease in cell wall autofluorescence in mature trichomes of *exo70H4* mutants as compared to WT plants. These deviations include also an absence of the callose-rich Ortmmanian ring (OR) from the *exo70H4* mutant trichomes (Kulich et al., 2015). The OR is a structure observed in mature trichomes that is rich in the β-1-3-glucan polymer callose and that divides trichomes into two major domains – the apical and the basal one (Kubátová et al., 2019; Kulich et al., 2018; Kulich et al., 2015). These domains are characterized by a different PM lipid composition and the presence of different EXO70 isoforms. EXO70A1 is recruited by the basal domain (located below the OR) and EXO70H4 is recruited by the apical domain, which is positioned above the OR (Kubátová et al., 2019). Deviations in cell wall autofluorescence and cell wall thickness connected to the defect in callose deposition were observed in the *exo70H4* mutant as compared to WT trichomes (Kulich et al., 2015). Only EXO70H4 but not the closely related EXO70H3 was able to rescue the aberrant callose deposition of *exo70H4* mutant trichomes, suggesting that EXO70H4 is crucial for a correct docking and PM secretion of callose synthases (Kulich et al., 2018).

Loss-of-function mutations in the *Mildew resistance locus o* (*Mlo*) gene of *Hordeum vulgare* (barley) have been known for decades to confer durable resistance against all naturally occurring isolates of the barley powdery mildew pathogen, *Blumeria hordei* (Jørgensen, 1992). The identification of barley *Mlo* (Büschges et al., 1997) revealed a medium-sized gene family (10–20 paralogues per plant species) encoding plant-specific and sequence-diversified integral membrane proteins (Devoto et al., 1999; Kusch et al., 2016). Members of this family have been identified in a plethora of mono- and dicotyledonous plant species (Devoto et al., 2003; Kusch et al., 2016), and natural or induced loss-of-function mutations of *Mlo* genes facilitated breeding for powdery mildew resistance in crop and legume species (reviewed in (Kusch and Panstruga, 2017)). In the dicotyledonous model plant *A. thaliana*, for instance, loss-of-function of three (*MLO2*, *MLO6*, *MLO12*) of the altogether 15 *MLO* genes (note the different nomenclature in barley and *A. thaliana*) is required to resemble the near complete resistance of barley *mlo* mutants (Consonni et al., 2006). In this scenario, the three proteins contribute unequally to the susceptibility of *A. thaliana* against the adapted powdery mildew pathogens *Golovinomyces orontii* and *G. cichoacearum*, with MLO2 (At1g11310) being the major contributor and MLO6 (At1g61560) as well as MLO12 (At2g39200) being minor contributors (Consonni et al., 2006). Apart from a reduced susceptibility to powdery mildew disease, barley *mlo* and *A. thaliana mlo2 mlo6 mlo12* plants develop indiscriminate callose-rich depositions at the cell wall and feature spontaneous death of mesophyll cells associated with a premature leaf senescence syndrome (Consonni et al., 2010; Piffanelli et al., 2002; Wolter et al., 1993). MLO2 further plays a decisive role in the regulation of ozone sensitivity (Cui et al., 2018) and in mediating systemic acquired resistance, a form of induced and systemically acting plant defense (Gruner et al., 2018).

Mutations in the *POWDERY MILDEW RESISTANT* (*PMR*) genes have been associated with a reduced susceptibility (i.e., enhanced resistance) of *A. thaliana* against these fungal pathogens (Vogel and Somerville, 2000). The respective forward genetics approach aimed for PR1-independent defense mechanisms and, accordingly, it was not surprising when *PMR2* was identified to be allelic to *MLO2* (Consonni et al., 2006). All other *PMR* genes uncloaked so far (*PMR4*–*PMR6*) encode proteins involved in the biogenesis and/or modification of the cell wall (Nishimura et al., 2003; Jacobs et al., 2003; Vogel et al., 2002; Vogel et al., 2004). PMR4 was shown to be a callose synthase (GSL5/CALS12) (Jacobs et al., 2003; Nishimura et al., 2003), PMR5 is a putative pectin O-acetyltransferase (Vogel et al., 2004; Chiniquy et al., 2019) and PMR6 a pectate lyase (Vogel et al., 2002). Consequently, MLO2 or other MLO proteins may be likewise involved in processes that are important for cell wall biogenesis.

MLO proteins were recently reported to function as calcium channels (Gao et al., 2022). Though this finding uncovered the long sought-after biochemical function of MLO proteins, it remains enigmatic how the calcium channel activity of MLO proteins connects to cellular processes and *mlo*-mediated powdery mildew susceptibility. Characteristic sequence features conserved among all MLO proteins are (i) a binding site for the ubiquitous calcium sensor calmodulin in the proximal part of the intracellular MLO carboxyl terminus (Kim et al., 2002), and (ii) four invariant cysteine residues in two of the extracellular loops, which form two disulfide bridges that are crucial for the stability of MLO proteins (Elliott et al., 2005). Resting upon phylogenetic analysis, MLOs were classified into seven clades while clade IV (mainly monocotyledonous plants) and clade V (mainly dicotyledonous plants) MLO proteins confer powdery mildew susceptibility (Kusch et al., 2016). Several studies carried out in *A. thaliana* reported that MLO proteins aside from clade V are involved in processes linked to polar secretion. For example, *A. thaliana mlo4* and *mlo11* mutant plants display severe root curling upon a tactile stimulus indicating a defect in thigmomorphogenesis (Chen et al., 2009; Bidzinski et al., 2014). Moreover, MLO7 (female gametophyte), as well as MLO5 and MLO9 (male gametophyte), are involved in the tête-à-tête between ovule and pollen tube as loss-of-function of these MLO proteins causes defects in ovule targeting by the pollen tube and, thus, reduced fertility (Kessler et al., 2010; Meng et al., 2020; Ju et al., 2021).

We here detected by a variety of histochemical staining procedures an unexpected phenotypic overlap of *A. thaliana exo70H4* and *mlo2 mlo6 mlo12* mutants regarding the biogenesis of trichome secondary cell walls. Prompted by this finding we combined genetic analysis with biochemical, cell biological protein-protein interaction and phytopathological assays, which together revealed an interplay of MLO and EXO70 proteins in the modulation of trichome cell wall composition and powdery mildew susceptibility by regulating secretory processes at the plasma membrane.

## Results

### Two independent *mlo2 mlo6 mlo12* triple mutants exhibit similar trichome cell wall phenotypes as the *exo70H4-1* mutant

While inspecting aniline blue-stained leaf specimens in the context of plant-powdery mildew pathoassays, we noticed that the rosette leaf trichomes of *mlo2-5 mlo6-2 mlo12-1* and *mlo2-6 mo-6-4 mlo12-8* triple mutants exhibited a delocalized callose phenotype that is reminiscent of the previously reported aberrant callose deposition pattern seen in trichomes of the exocyst subunit *exo70H4-1* mutant (Kulich et al., 2018). To assess this feature quantitatively, we defined three distinctive callose phenotype categories (**Figure 1**) and determined the percentage of trichomes belonging to the respective category for each mutant genotype, including a corresponding Col-0 WT control (**Figure 1A**). We found that ∼68% of trichomes in Col- 0 showed a prototypical WT-like phenotype, which is characterized by aniline blue-based fluorescence of the OR region with or without additional fluorescence of the basal regions of the trichome branches. In addition, ∼14% of the WT trichomes exhibited a delocalized callose pattern and ∼17% lacked any detectable fluorescence (callose deposition), possibly due to the different developmental stages of the inspected trichomes (**Figure 1B**). By contrast, the *exo70H4-1* mutant exhibited a drastically perturbed trichome staining pattern (< 1% WT-like OR, ∼28% delocalized, and ∼72% no callose deposition, respectively) as reported previously (Kulich et al., 2015 and 2018). This phenotypic deviation was shared by the two *mlo* triple mutants, which showed a similar reduction in the WT-like fluorescence pattern as well as concomitant increases in aberrant callose deposits and an apparent lack of callose deposition (*mlo2-5 mlo6-2 mlo12-1*: ∼2% WT-like OR, ∼48% delocalized and ∼50% no callose deposition, respectively; *mlo2-6 mlo-6-4 mlo12-8*: **∼**7% WT-like OR, ∼32% delocalized and ∼61% no callose deposition, respectively) (**Figure 1B**).

**Figure 1.**
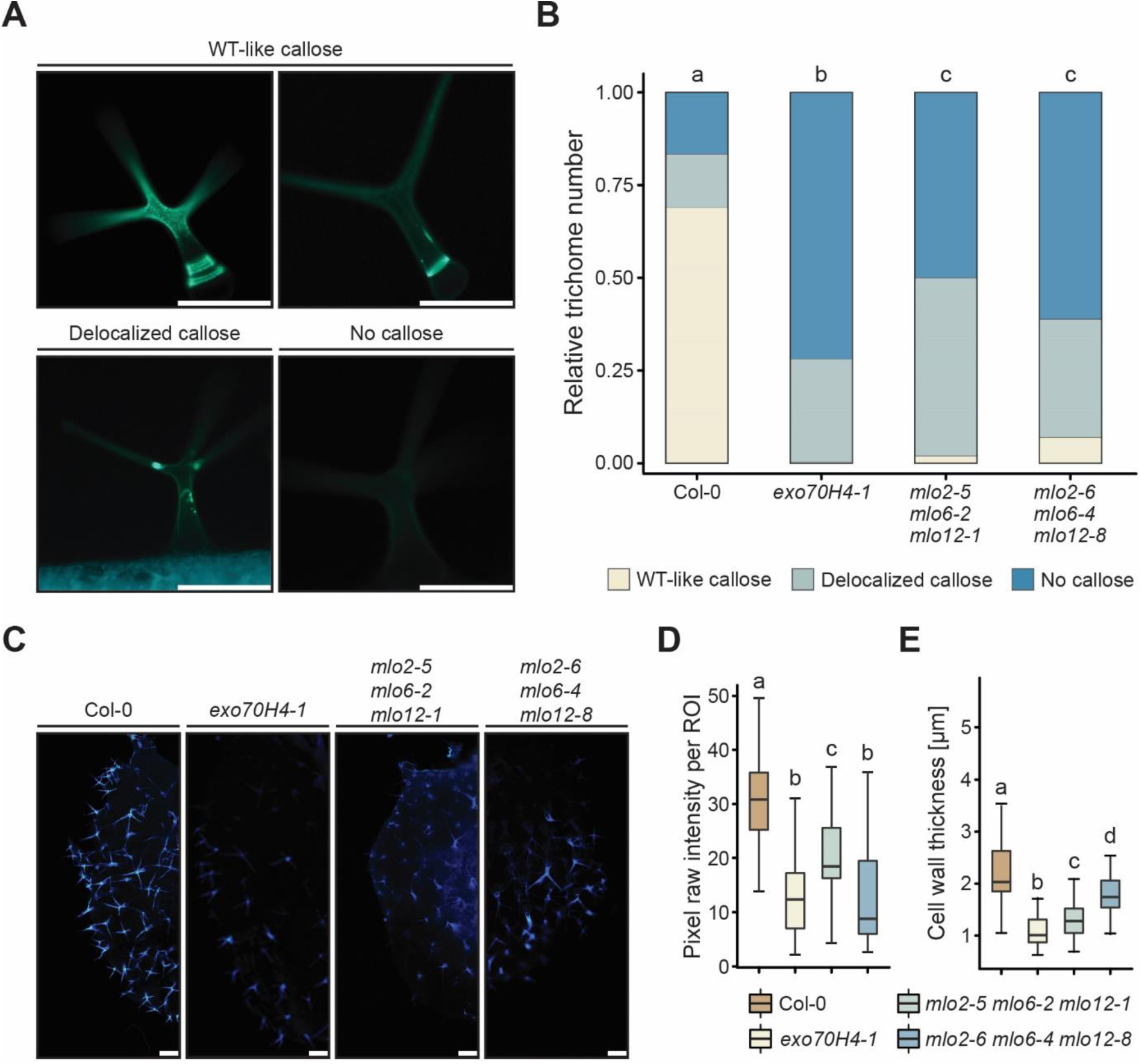
The *exo70H4-1* mutant and two independent *mlo2 mlo6 mlo12* triple mutants share similar rosette leaf trichome phenotypes. **A**) Representative micrographs illustrating trichome-associated callose phenotype categories (WT-like callose, delocalized callose, no callose) upon staining with aniline blue. **B**) Distribution of trichome callose patterns in trichomes of Col-0 WT and the indicated mutants upon staining with aniline blue. Values are based on four experiments with about 20 trichomes inspected per experiment and genotype. Data were analyzed by chi-square test, the *p*-values were corrected by FDR (*α* = 0.05), and letters denote statistically significant differences between genotypes. **C**) Representative micrographs depicting trichome cell wall autofluorescence of Col-0 WT plants and the indicated mutants. **D**) Quantification of trichome cell wall autofluorescence. Shown is the pixel intensity per region of interest (ROI), calculated based on autofluorescence micrographs as those exemplarily shown in (**C**). Values are based on two experiments with about 20 trichomes inspected per experiment and genotype. Letters assign differences of statistical significance (pairwise Wilcoxon-Mann-Whitney test corrected by FDR, *α* = 0.01). **E**) Quantification of trichome cell wall thickness. Cell wall thickness was measured in proximity to the OR. Values are based on four experiments with 20 trichomes inspected per experiment and genotype. Letters assign differences of statistical significance (pairwise Wilcoxon-Mann-Whitney test corrected by FDR, *α* = 0.01). Scale bars in (**A**) and (**C**) represent 200 µm.

Inspired by the intriguing parallels in trichome-associated callose deposition phenotypes between the *exo70H4-1* mutant and the two *mlo* triple mutants, we explored whether the trichomes of these mutants exhibit further commonalities. To this end, we next measured cell wall autofluorescence (**Figure 1C** and **D**) and cell wall thickness (**Figure 1E**). We observed that the trichomes of the *exo70H4-1* mutant and the two *mlo2 mlo6 mlo12* triple mutants showed significantly less autofluorescence than those of Col-0 WT plants (**Figure 1C** and **D**). Similarly, we found that the trichomes of all three investigated mutant genotypes developed significantly thinner cell walls than Col-0 WT trichomes (**Figure 1E**).

We next subjected rosette leaves of the above-described WT and mutant genotypes to histochemical staining to study the *in situ* accumulation of reactive oxygen species (ROS) and heavy metals (HMs) in leaf trichomes. We chose these two features since trichomes of the *exo70H4-1* mutant were previously found to have defects in the accumulation of ROS and HMs (Kulich et al., 2015). As described above for the callose staining pattern (**Figure 1A** and **B**), we defined phenotypic categories and determined the proportion of trichomes in the respective category for each histochemical dye and genotype (**Figure 2**). Upon staining the specimens with 3,3’-diaminobenzidine (DAB) to visualize ROS accumulation, we observed that the majority (∼53%) of Col-0 WT trichomes exhibited a demarcated region of ROS localization in the basal part of the trichomes, coinciding with the presumed location of the OR. A substantial portion (∼43%) of Col-0 trichomes also lacked detectable ROS accumulation (though sometimes weak staining was seen in branch tips), while only a minority (∼3%) showed a delocalized ROS pattern, which was characterized by a broader dispersal extending into the branching regions of the trichomes (**Figure 2A** and **B**). By contrast, *exo70H4- 1* mutant plants largely lacked any ROS accumulation in the trichomes or displayed an aberrant ROS distribution (∼59% and ∼39%, respectively. The same was essentially true for the two *mlo2 mlo6 mlo12* triple mutants, which like the *exo70H4-1* mutant showed a phenotypic pattern that differed from Col-0 WT trichomes in a statistically significant manner (**Figure 2A** and **B**).

**Figure 2.**
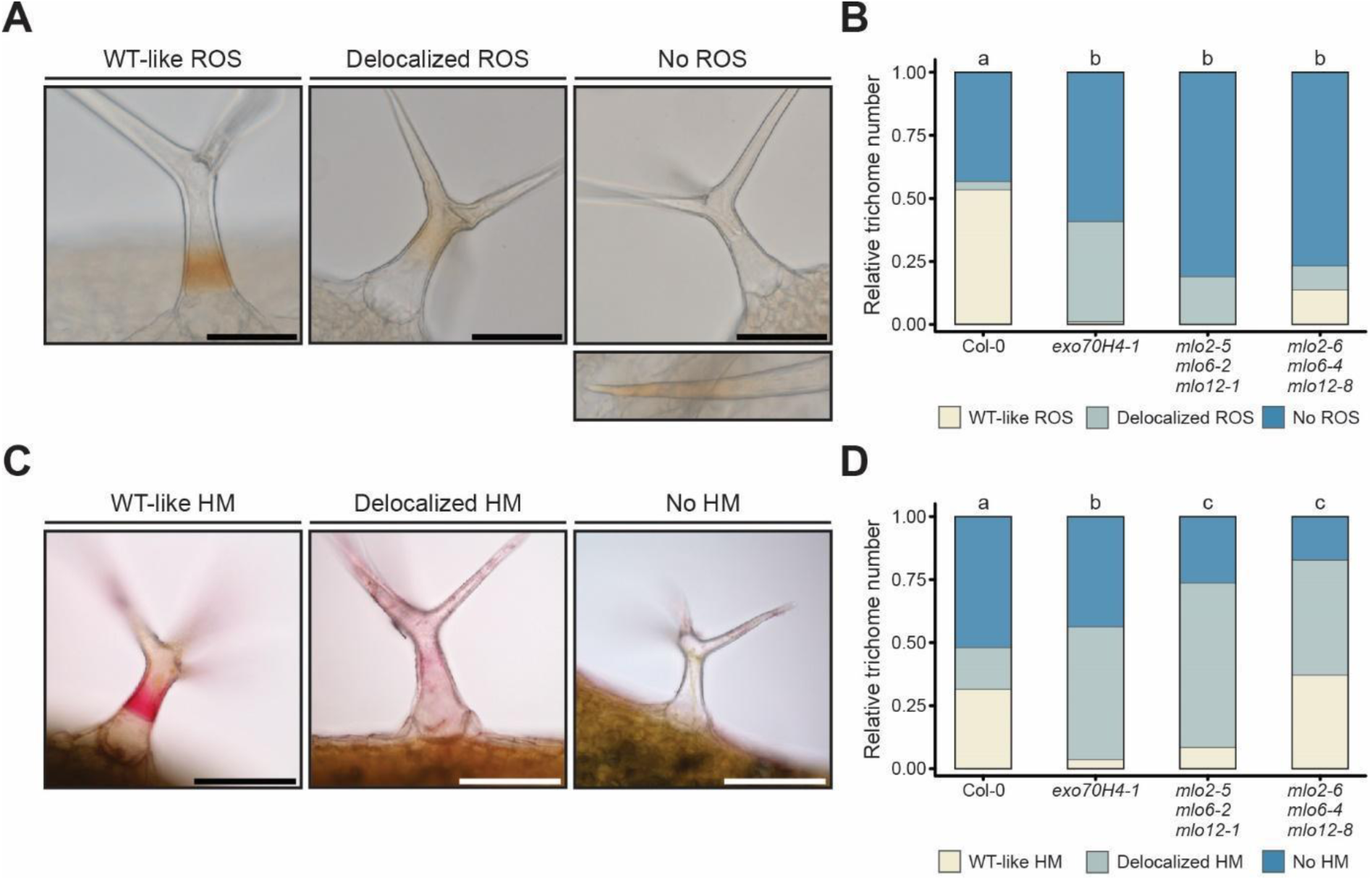
The *exo70H4-1* single mutant and the *mlo2 mlo6 mlo12* triple mutants share similar aberrant ROS and HM localization patterns in rosette leaf trichomes. A) Representative micrographs illustrating trichome-associated reactive oxygen species (ROS) phenotype categories (WT-like, delocalized ROS, no ROS). Trichomes lacking ROS accumulation in the basal part of the trichomes and the proximal areas of the branches occasionally exhibited weak DAB staining in the branch tips (see separate micrograph below the main panel). **B**) Distribution of trichome ROS patterns in Col-0 WT and the indicated mutants. Values are based on three experiments with approximately 50 trichomes inspected per experiment and genotype. Data were analyzed by chi-square test, the *p*-values were corrected by FDR (*α* = 0.05), and letters denote statistically significant differences between genotypes. **C**) Representative micrographs illustrating trichome-associated heavy metal (HM) phenotype categories (WT-like, delocalized, no HM). **D**) Distribution of trichome HM patterns in Col-0 WT and the indicated mutants. Values are based on three experiments with approximately 50 trichomes inspected per experiment and genotype. Data were analyzed by chi-square test, the *p*-values were corrected by FDR (*α* = 0.05), and letters denote statistically significant differences between genotypes. Scale bars in (**A**) and (**C**) represent 200 µm.

The histochemical analysis of HM deposition by diphenylthiocarbazone (dithizone) staining revealed similar deviations as the DAB staining for ROS accumulation described above. Leaf trichomes of Col-0 WT plants showed either ring-like staining of the basal trichome regions (∼30%) or no staining (∼50%), and only rarely exhibited delocalized staining (∼16%). Differing from this distribution, trichomes of the *exo70H4- 1* single mutant and the two *mlo2 mlo6 mlo12* triple mutants had a greater proportion of delocalized staining (>40%), which in all cases was different from Col-0 WT in a statistically significant manner (**Figure 2C** and **D**). Collectively, these assays revealed an unexpected phenotypic overlap between the *exo70H4-1* mutant and *mlo2 mlo6 mlo12* triple mutants that points to shared alterations in trichome cell wall architecture of these mutants and therefore to a common/shared function of the associated proteins in trichome cell wall biogenesis.

### Genetic analysis identifies mutations in *MLO2* and *MLO6* as the main cause of the aberrant trichome phenotypes in the *mlo* triple mutants

To assess whether the trichome phenotypic deviations observed with the two *mlo2 mlo6 mlo12* triple mutants were due to one of the mutated *MLO* genes or the effect of two or all three mutations in combination, we performed some of the above-described assays also with a selected subset of *mlo2*, *mlo6,* and *mlo12* single and double mutants. Regarding callose staining with aniline blue, we observed that the *mlo6*-*2* single mutant, as well as the *mlo2-5 mlo6-2* and *mlo6-2 mlo12-1* double mutants, showed a distribution pattern that largely resembled the pattern of the two *mlo* triple mutants concerning the occurrence of delocalized callose deposits or the absence of callose deposition. By contrast, the *mlo2-5* and *mlo12-1* single mutants as well as the *mlo2-5 mlo12-1* double mutant had a WT-like distribution of callose staining (**Supplemental Figure 1A**). A similar tendency, yet less pronounced, was visible for the thickness of trichome cell walls, with the *mlo6-2* mutant and its derived double mutants showing the largest deviation from Col-0 WT trichomes (**Supplemental Figure 1B**). Concerning ROS accumulation, trichomes of all *mlo* single and double mutants were significantly different from Col-0 WT trichomes (**Supplemental Figure 1C**). Regarding trichome autofluorescence, all *mlo* single and double mutants apart from the *mlo2-5 mlo6-2* double mutant, which showed drastically reduced autofluorescence, exhibited a WT-like fluorescence intensity (**Supplemental Figure 1D**). Finally, with respect to the accumulation of HMs in trichomes, particularly trichomes of the *mlo2-5* single mutant and the *mlo2-5 mlo6-2* double mutant revealed distribution patterns that deviated from Col-0 WT and resembled the phenotype of *mlo* triple mutant plants (**Supplemental Figure 1E**). In sum, we noticed a variable contribution of the three *MLO* genes to the various trichome phenotypes. Nonetheless, a pivotal contribution of *MLO6*, alone or in combination with *MLO2*, emerged as a common theme across the different assays. However, due to the mixed and variable contribution of the three *MLO* genes to the trichome phenotypes, we kept performing the following experiments with the two *mlo2 mlo6 mlo12* triple mutants for consistency.

### Cell wall analysis of isolated trichomes provides evidence for altered cell wall composition in the *exo70H4-1* single mutant and the two *mlo* triple mutants

We next wondered whether the similar aberrations in *exo70H4-1* and *mlo2 mlo6 mlo12* mutant trichomes were symptoms of pervasive alterations of cell wall composition in these trichomes compared to the Col-0 WT. Intending to subject isolated leaf trichomes to global quantitative cell wall analyses, we first used histochemistry to validate that the cell wall deviations observed in *exo70H4-1* and *mlo2 mlo6 mlo12* leaf- resident trichomes are likewise detectable in detached trichomes (**Figure 3**). We isolated trichomes *via* an established method for the gentle separation of rosette leaf tissue and trichomes (Huebbers et al., 2022) and stained the isolated trichomes with calcofluor white, which binds to unsubstituted β-glucan polymers such as cellulose or callose (Eschrich and Currier, 1964; Wood, 1980). A quantitative microscopic assessment revealed that ∼64% of the Col-0 trichomes showed an OR-like structure in their basal regions (**Figure 3A**). By contrast, trichomes retrieved from *exo70H4-1* and *mlo2-5 mlo6-2 mlo12-1* mutant plants displayed a statistically significant reduction in OR formation to ∼10% and ∼7% of the inspected trichomes, respectively. In contrast to leaf-attached trichomes (**Figure 1A**), OR formation in isolated trichomes of *mlo2-6 mlo6-4 mlo12-8* mutant plants (∼52%) resembled WT-like numbers, possibly due to differences in dye uptake between leaf-associated and detached trichomes. In sum, the calcofluor white staining pattern of isolated trichomes was largely consistent with the aniline blue staining pattern of leaf-associated trichomes described above (**Figure 1A** and **B**).

**Figure 3.**
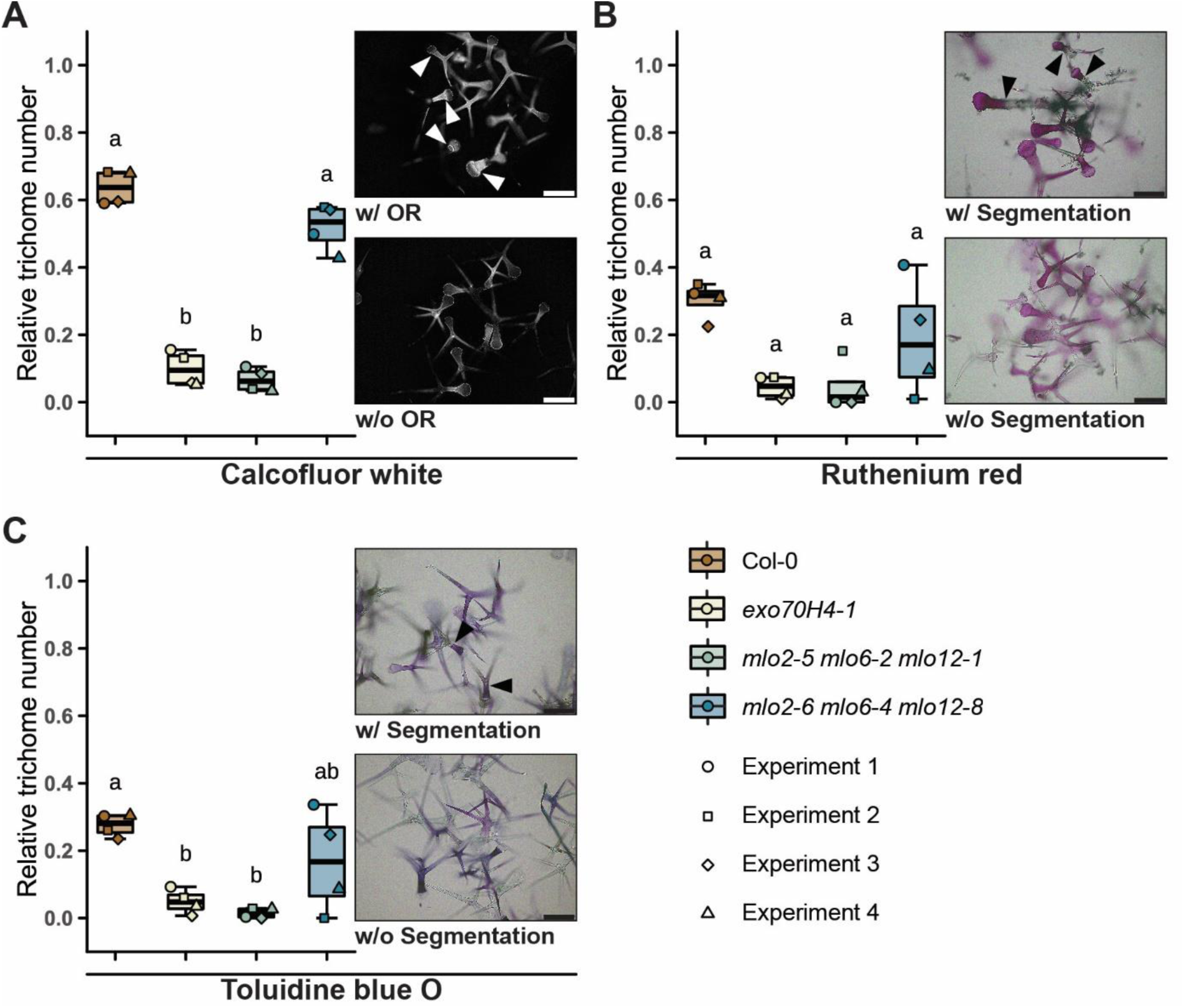
Histochemical staining of detached rosette leaf trichomes indicates a reduced frequency of cell wall segmentation in *exo70H4-1* and *mlo2 mlo6 mlo12* mutants. Rosette leaves trichomes were isolated and stained as described in the Materials and Methods section. **A**) Frequency of the OR upon calcofluor white staining. **B** and **C**) Frequency of basal-apical segmentation upon ruthenium red (**B**) or toluidine blue O (**C**) staining. Boxplots indicate the outcome of four independent experiments with three replicates each. About 100 trichomes were assessed per replicate, i.e. approx. 1,200 trichomes per genotype in total. Letters assign differences of statistical significance (pairwise Student’s *t*-test corrected by FDR, *α* = 0.01). Micrographs show stained trichomes with (w/, arrowheads) or without (w/o) the respective pattern. Scale bars represent 200 µm.

We also subjected the isolated trichomes to the colorimetric dyes ruthenium red (visualization of pectin) and toluidine blue O. The latter has been described to stain polygalacturonic acids pinkish-purple and poly-aromatic substances such as lignin blue or greenish-blue (Pradhan Mitra and Loqué, 2014). Though these stains did not yield hints for drastic differences in pectin or lignin content across the various genotypes, we encountered a striking segmentation pattern in ∼30% of the Col-0 trichomes on average for ruthenium red and toluidine blue O (**Figure 3B** and **C**). This pattern was characterized by a stained basal and an unstained apical trichome region, separated by a sharp boundary that differed in its position from the location of the OR. Unlike for WT trichomes, the formation of this clear segmentation was considerably reduced in *exo70H4-1*- (∼5%) and *mlo2-5 mlo6-2 mlo12-1* (∼3%)-derived trichomes. Trichomes released from *mlo2-6 mlo6-4 mlo12-8* mutant plants, by contrast, occasionally (∼17%) exhibited a WT-like segmentation pattern, accompanied by a higher degree of variation across the experiments (**Figure 3B** and **C**).

We next subjected isolated WT and mutant trichomes to analytical quantification of cell wall matrix monosaccharides (rhamnose, arabinose, galactose, mannose, fucose, xylose, and glucose), uronic sugar acids, and crystalline cellulose (**Figure 4** and **Supplemental Figure 2**). For most of the neutral monosaccharides analyzed, there was no statistically significant difference in their levels between the mutants and Col-0 WT trichomes (**Supplemental Figure 2A–E**). Exceptions were arabinose and galactose, for which lower levels were found in trichomes of either the *mlo2-5 mlo6-2 mlo12-8* mutant (arabinose; **Figure 4A**) or both *mlo* triple mutants (galactose; **Figure 4B**). Taken together, these deviations caused a reduction in the sum of the neutral monosaccharides in the case of the *mlo2-5 mlo6-2 mlo12-8* mutant as compared to the other genotypes (**Figure 4C**). Similar to most neutral monosaccharides, we also did not find any marked differences in crystalline cellulose content and the levels of uronic acids between mutant and Col-0 WT trichomes (**Supplemental Figure 2F and G**).

**Figure 4.**
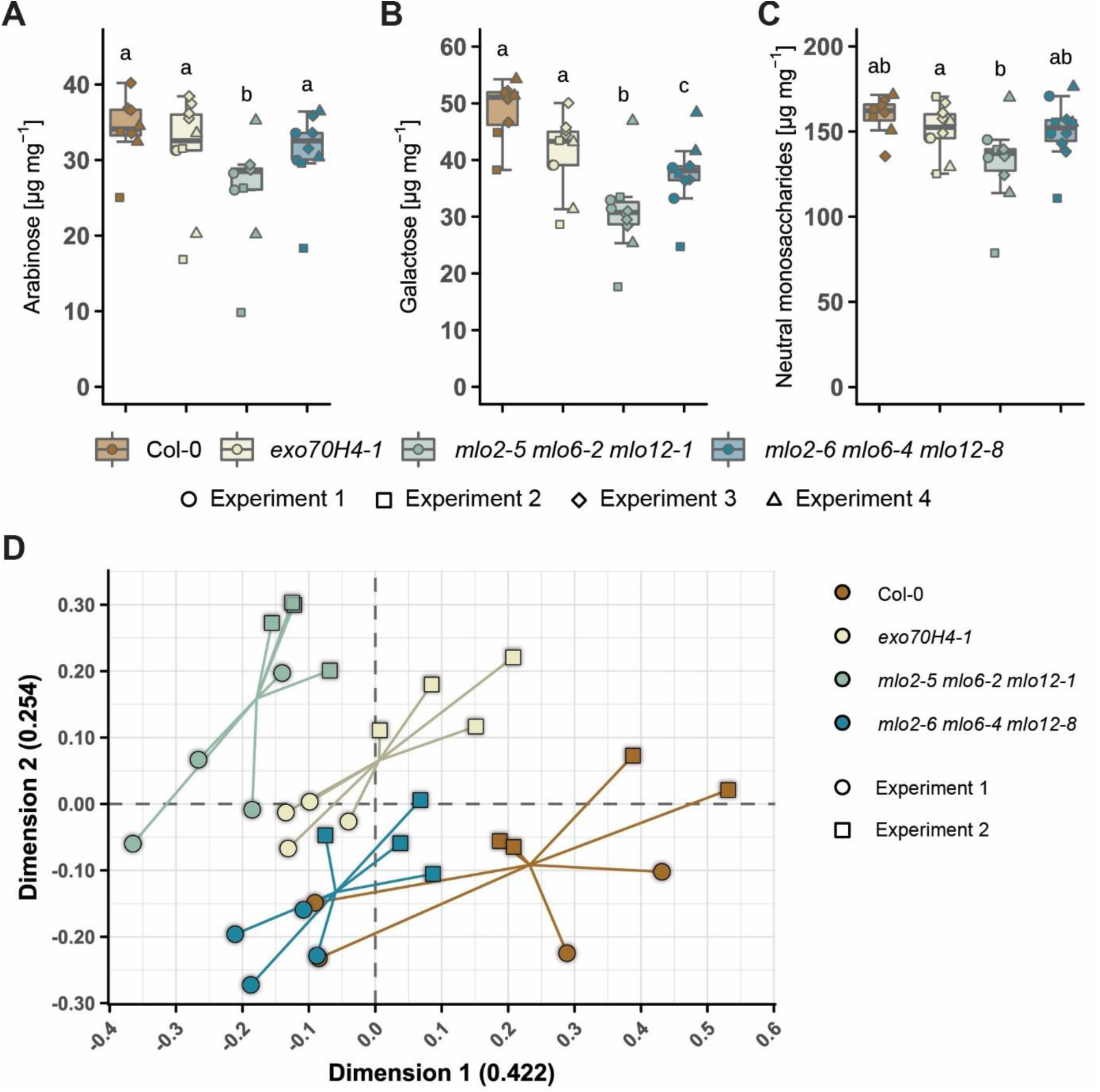
Cell walls of isolated *exo70H4-1* and *mlo2 mlo6 mlo12* triple mutant rosette leaf trichomes have altered carbohydrate composition. A-C) Quantification of neutral matrix monosaccharides. Genotypes are indicated by different colors whereas different geometric shapes denote samples that arose from the same independent experiment, as indicated in the legend below the boxplots. **A** and **B**) Abundance of the neutral monosaccharides arabinose (**A**) and galactose (**B**) in the alcohol-insoluble residue recovered from trichomes of the various genotypes. The boxplot indicates the outcome of at least three independent experiments with two to four replicates each. Letters assign differences of statistical significance (pairwise Student’s *t*-test corrected by FDR, *α* = 0.01). **C**) Total abundance of neutral monosaccharides in the cell wall matrix of the various genotypes. The boxplot includes the amounts of the monosaccharides indicated in panels (**A**) and (**B**) in addition to the abundance of rhamnose, mannose, fucose, xylose, and matrix glucose in the alcohol- insoluble residue (**Supplemental** Figure 2A**–E**). Letters assign differences of statistical significance (pairwise Student’s *t*-test corrected by FDR, *α* = 0.01). **D**) Principal component analysis of FTIR spectra of trichomes isolated from the various genotypes. The plot shows the first and second dimensions, which accounted for 42.2% and 25.4% of the variability between samples, respectively. The intersection point of the lines indicates the mean of the data points per genotype.

We subjected the same sample material used for the quantification of uronic acids to Fourier transform infrared (FTIR) spectroscopy for the global analysis of cell wall composition. Averaging of the retrieved spectra per genotype and visualization in a plot revealed notable differences between the genotypes tested (**Supplemental Figure 2H**). Areas that stood out in this averaged plot included (i) a peak (indicating a reduction in comparison to Col-0) in the region of ∼1,500 cm^-1^ to ∼1,300 cm^-1^ prominent in all mutants, (ii) a peak between ∼1,200 cm^−1^ and ∼900 cm^-1^ in both *mlo* triple mutants accompanied by a high variability as indicated by the shaded area, and (iii) a peak around 850 cm^-1^ in all mutants. The FTIR spectra of the individual measurements from two independent experiments (eight samples per genotype in total) were used for principal component analysis (PCA). We assembled a plot of the first and second dimension, which accounted for 42.2% and 25.4% of the variability within the data, respectively (**Figure 4D**). The PCA plot indicates an overall genotype-specific clustering of the individual spectra, albeit two spectra of Col-0 trichomes grouped with the spectra retrieved from *mlo2-6 mlo6-4 mlo12-8* samples. Despite this minor deviation, we concluded that all mutant samples differ from the WT in dimension one, whereas only *exo70H4-1* and *mlo2-5 mlo6-2 mlo12-1* samples, but not *mlo2-6 mlo6-4 mlo12-8* samples, vary also in dimension two. The eigenvectors associated with such wavenumbers that contributed the most to dimensions one and two are listed in **Supplemental Table 1** and comprised values between 1,474 cm^-1^ and 1,425 cm^-1^ as well as values around 900 cm^-1^, which are linked to various cell wall constituents such as cellulose, pectin, and arabinogalactan. Altogether, the results of FTIR spectroscopy indicated global alterations in the cell walls of *exo70H4-1* single and *mlo* triple mutant trichomes.

### Fluorophore-labeled EXO70H4 and MLO proteins co-localize in leaf trichomes

To study the subcellular localization of EXO70H4 and MLO proteins in rosette leaf trichomes, we transformed mCherry-EXO70H4-expressing plants in the Col-0-derived *rdr6* background (Kulich et al. 2018), which reduces transgene silencing, with constructs expressing GFP-tagged versions of MLO2, MLO6, and MLO12, respectively. All *MLO* genes were expressed under the control of the *EXO70H4* promoter, which confers preferential expression in trichomes (Kulich et al., 2018). Inspection of the resulting transgenic lines by confocal laser scanning microscopy (**Figure 5**) revealed that the MLO2-GFP fusion protein co-localized with mCherry- EXO70H4 in trichomes, mainly in the zone of the OR and above it at the plasma membrane (PM) delineating the thickened secondary cell wall. Occasionally we also observed co-localization of the two proteins in the apical domain of the trichome PM. Here, MLO2-GFP co-localized with mCherry-EXO70H4 in PM speckles and the cell wall (**Figure 5A**). Similar subcellular distribution patterns were observed for the MLO6- GFP and MLO12-GFP fusion proteins, which co-localized with mCherry-EXO70H4 at the OR region, in PM speckles and the cell wall (**Figure 5A**). Interestingly, the domain defined by MLO12-GFP appeared to be broader than the mCherry-EXO70H4 signal, which localized to its center, as if delimited by the MLO12-GFP protein as a boundary (**Supplemental Figure 3)**. Furthermore, we detected mobile MLO12-GFP-labeled bodies in the cytoplasm. To our surprise, we observed both EXO70H4 and all three fluorescent MLO fusion proteins also in cell wall channels, which were oriented perpendicularly to the PM and appeared to be connected to the cytoplasm (**Figure 5A**). This observation indicates the trapping of fluorophore-labeled EXO70H4 and MLO12 within the cell wall – a conclusion supported by the visualization of membrane remnants within the cell wall as visualized by electron microscopy (Kubátová et al., 2019). Altogether, these data suggest that mCherry-EXO70H4 co-localizes with all three GFP-labeled MLO proteins tested, further highlighting the possibility that they might operate in the same cellular pathway and may even co-reside within the same macromolecular complex/compartment.

**Figure 5.**
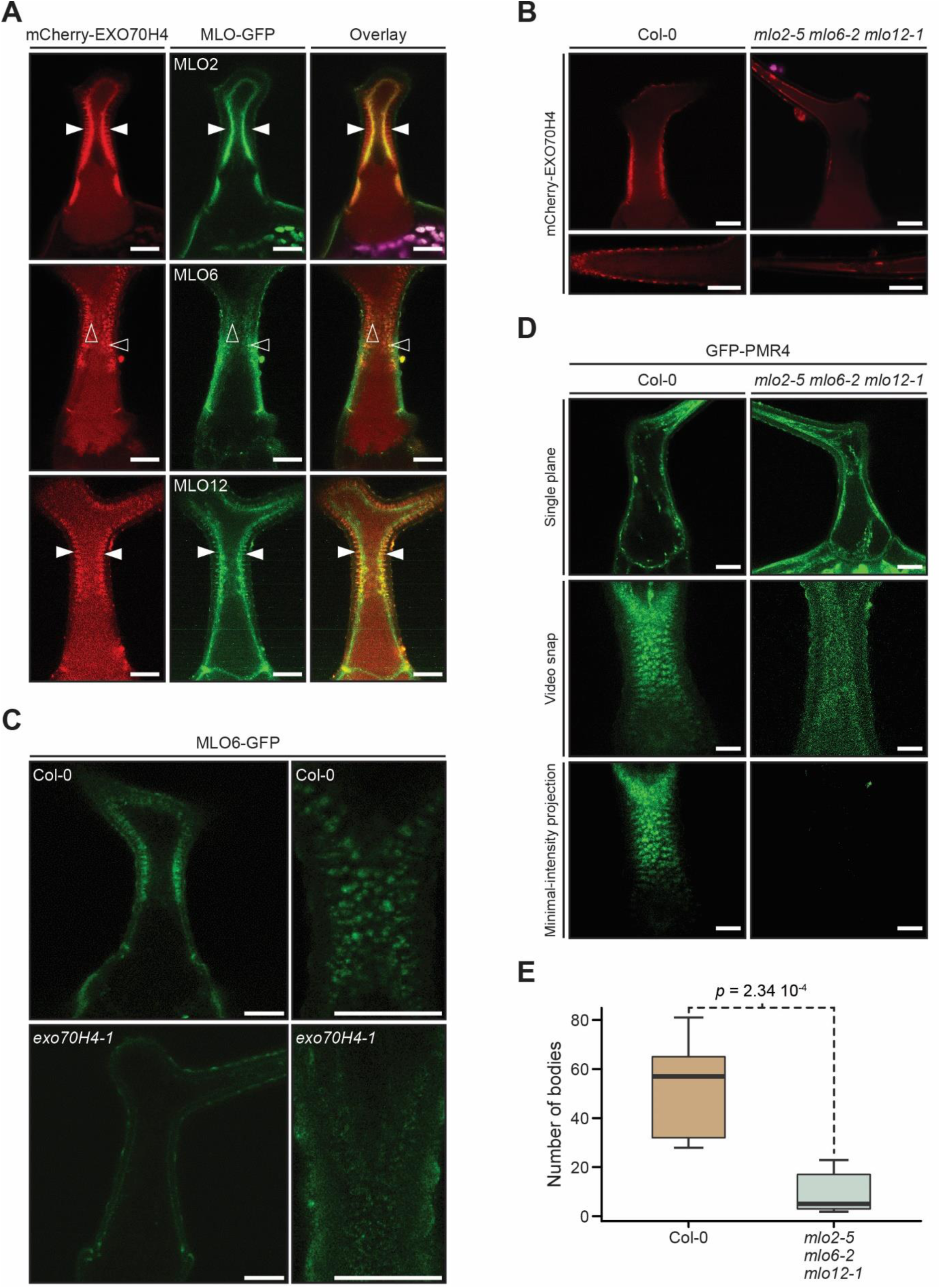
Subcellular localization of fluorophore-labeled EXO70H4, MLO, and PMR4 proteins in rosette leaf trichomes of *exo70H4* and *mlo2 mlo6 mlo12* transgenic lines. A) Co-localization of mCherry-EXO70H4 and GFP-tagged MLO proteins in trichomes of transgenic *A. thaliana* lines. Both mCherry-EXO70H4 and MLO-GFP proteins (MLO2, MLO6, and MLO12) were expressed in transgenic *A. thaliana* lines (Col-0-derived *rdr6* background) under the control of the *EXO70H4* promoter. In each micrograph, a single confocal plane is shown. Filled and open arrowheads point to examples of the fluorophore-labeled cell wall channels and PM speckles, respectively. Scale bars represent 20 μm. **B**) Subcellular localization of mCherry-EXO70H4, expressed under the control of the *EXO70H4* promoter in either a transgenic Col-0 WT plant (left panels) or a *mlo2-5 mlo6-2 mlo12-1* triple mutant plant (right panels). The upper panels show the trichome stalk region, the lower panels depict an individual branch. Scale bars represent 20 μm. **C**) Subcellular localization of MLO6-GFP, expressed under the control of the *MLO6* promoter, in either a transgenic Col-0 WT plant (upper panels) or the *exo70H4-1* mutant plant (lower panels). The left panels show single planes focused on the cell wall, the right panels show single planes focused on the plasma membrane. Scale bars are 20 μm each. **D**) Subcellular localization of GFP-PMR4, expressed under the control of the *EXO70H4* promoter in either a transgenic Col-0 WT plant (left panels) or a *mlo2-5 mlo6-2 mlo12-1* triple mutant plant (right panels). The micrographs show a single confocal plane (upper panels), a snapshot from a time-series video (middle panels), and a minimal-intensity projection from the time series (bottom panels). **E**) Quantification of the number of stationary GFP-PMR4-labeled intracellular dot-like compartments as calculated from the minimal-intensity projection images (see Materials and Methods for further details). Data are based on two experiments with a total of 11 (Col-0) and 21 (*mlo2-5 mlo6-2 mlo12-1* triple mutant) trichomes analyzed. The *p*-value was computed according to Wilcoxon-Mann-Whitney and indicates the statistical difference between the mobility of GFP-PMR4 in the Col-0 WT background and the *mlo2−5 mlo6−2 mlo12−1* background.

### EXO7H4 and MLO proteins mutually impact each other’s subcellular localization

Since EXO70H4 and MLO proteins co-localize substantially in trichomes, we next investigated whether they influence each other’s localization. We transformed Col-0 WT plants and the *mlo2-5 mlo6-2 mlo12-1* mutant with the above-mentioned mCherry- EXO70H4 construct and comparatively analyzed the subcellular localization of the fluorescent fusion protein in the resulting transformants. In transgenic Col-0 plants, we observed that the mCherry-EXO70H4 signal is mostly present at the PM and within the thickened secondary cell wall of the central trichome stalk region (**Figure 5B**). In addition, red fluorescence was localized at the cell wall papillae that decorate the surface of individual trichome branches. By contrast, mCherry-EXO70H4 exhibited predominantly cytoplasmic localization in *mlo2-5 mlo6-2 mlo12-1* transformants and was essentially absent from the PM and the cell wall, including its surface papillae (**Figure 5B**).

As the *mlo6-2* mutant most strongly resembled the aberrant trichome phenotypes of the *exo70H4-1* mutant (**Supplemental Figure 1**), we focused on MLO6 to study whether the *exo70H4* mutation likewise affects the subcellular localization of MLO proteins. To this end, stably expressed the MLO6-GFP fusion protein under the control of the native *MLO6* promoter in both, Col-0 WT and *exo70H4-1* mutant plants. We noticed that in the trichomes of transgenic Col-0 WT plants, MLO6-GFP seemed to localize primarily to the PM and the cell wall, including the above-described cell wall channels (**Figure 5C**), which is reminiscent of its co-expression with mCherry- EXO70H4 in the *rdr6-1* mutant background (**Figure 5A**). In addition, we detected the MLO6-GFP fusion protein in stationary PM-associated speckles (**Figure 5C**). By contrast, the MLO6-GFP fluorescent signal was less pronounced in the cell walls of *exo70H4-1* transgenic plants, possibly due to the reduced thickness of trichome secondary cell walls in this genetic background (Kulich et al., 2015). Instead, MLO6- GFP was in part visible in the cytoplasm in this genotype. In addition, the PM-associated MLO6-GFP speckles were smaller and appeared more mobile in the *exo70H4-1* transgenic lines. In summary, the data suggest that EXO7H4 and MLO proteins mutually impact each other’s subcellular localization.

### Reminiscent of transgenic *exo70H4-1* lines, PMR4 callose synthase is mislocalized in transgenic *mlo2-5 mlo6-2 mlo12-1* triple mutant lines

Because of the similarities in aberrant callose deposition in the *exo70H4* and *mlo2-5 mlo6-2 mlo12-1* triple mutant (see above and **Figure 1**), we decided to examine the subcellular localization and intracellular dynamics of the PMR4 callose synthase in the *mlo* triple mutant. PMR4 is responsible for callose deposition in *A. thaliana* trichomes and delocalizes in the *exo70H4-1* mutant (Kulich et al., 2018). Interestingly, when we expressed a GFP-tagged version of PMR4 in stably transformed *mlo2-5 mlo6-2 mlo12-1* triple mutant plants, we did not observe the characteristic cell wall localization pattern of the protein seen in PMR4-transgenic Col-0 WT plants. Instead, GFP-PMR4 showed higher intracellular abundance in the *mlo* triple mutant, as revealed by the imaging of single confocal planes (**Figure 5D**). Moreover, the number of clustered immobile GFP-PMR4-labeled endomembrane bodies (speckles), which likely represent PMR4 compartments formed during exocytosis (Kulich et al., 2018), was significantly lower in the PMR4-transgenic *mlo2-5 mlo6-2 mlo12-1* triple mutant plants than in the respective transgenic Col-0 plants, as revealed by time series imaging and minimal-intensity projection microscopy (**Figure 5D and E**). The latter technique enables to discriminate mobile from stationary structures in time-lapse series. This data indicates that the delivery of PMR4 to the plasma membrane and, thus, ultimately to the secondary cell wall, is perturbed in the *mlo* triple mutant, similarly as previously observed for the *exo70H4-1* mutant (Kulich et al., 2018).

### Clade V MLO proteins interact with different EXO70 proteins in an isoform-preferential manner

Based on the similar phenotypic aberrations of rosette leaf trichomes in the *exo70H4- 1* mutant and *mlo2 mlo6 mlo12* triple mutants (**Figure 1** and **2**) and the co-localization of EXO70H4 and MLO proteins in trichomes (**Figure 5A**), we hypothesized that EXO70H4 and MLO2, MLO6 and MLO12 might not only be active in the same cellular pathway but may also physically interact. Furthermore, we speculated that such an interaction would not inevitably be confined to EXO70H4 but may also involve other EXO70 isoforms. As gene co-expression can serve as an indicator of co-functionality in a particular process (Humphry et al., 2010; Gupta and Pereira, 2019), we took advantage of published transcript data (Winter et al. 2007) to identify potential candidate EXO70 proteins for an interaction screening with MLO proteins (**Figure 6**). Using tissue-specific or biotic stress-associated data sets, we carried out a co-expression analysis and calculated Pearson correlation coefficients (*ρ*) for all combinations of *MLO* and *EXO70* genes in *A. thaliana* (**Figure 6A**, note that no transcript data were available for *EXO70A1* and -*H6*). We placed particular emphasis on *EXO70* genes that showed a high and positive correlation with genes encoding clade V MLO proteins (MLO2, MLO6, and MLO12). With regards to tissue-specific data, we found that *MLO2* expression was correlated with *EXO70D2* (*ρ* = 0.63) and *EXO70E1* (*ρ* = 0.62) (**Figure 6A**, Tissue-specific). Strikingly, *MLO6* showed a high correlation with *EXO70H4* (*ρ* = 0.63), consistent with the notion that both genes were found to be strongly expressed in trichomes (Marks et al., 2009; Gilding and Marks, 2010). Concerning biotic stress-associated expression, *MLO2*, *MLO6,* and *MLO12* but also the closely related *MLO3* displayed strongly correlated expression with the *EXO70* genes *-B1* and *-B2*, *-D3*, *-E1* and *-E2*, *-F1* as well as -*H1* and *-H2* (**Figure 6A**, Biotic stress). Especially the expression of *EXO70H1* and *EXO70H2* showed a high level of correlation with *MLO12* (*ρ* = 0.93).

**Figure 6.**
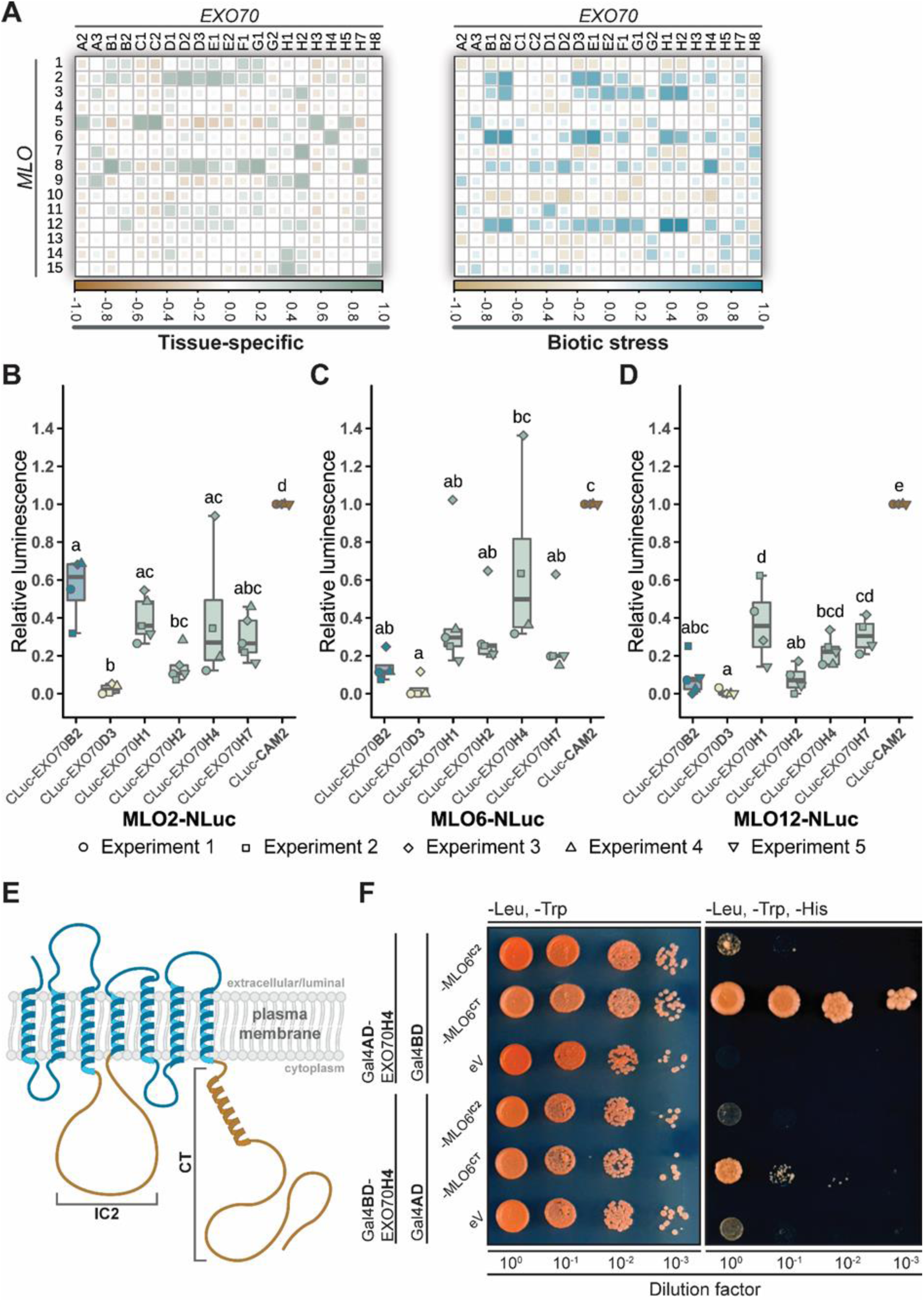
Interaction between clade V MLO proteins and different EXO70 proteins. **A**) Tissue-specific or biotic stress-induced correlation of *MLO* and *EXO70* transcript abundance. Transcript data (note that no data were available for *EXO70A1* and *EXO70H6*) were retrieved from the Arabidopsis eFP browser (http://bar.utoronto.ca/efp/cgi-bin/efpWeb.cgi) on 2021-04-19. Pearson correlation was calculated to assess the co-expression of individual *MLO* and *EXO70* pairs. Squares illustrate Pearson correlation coefficients as indicated by the color code below each graph. **B–D**) Luciferase complementation of different CLuc-EXO70 and MLO- NLuc protein variant combinations was carried out by transient gene expression in *N. benthamiana*. Representative leaves exhibiting luminescence signals detected for the various combinations are shown in **Supplemental** Figure 5A. For normalization, luminescence signals were divided by the signal obtained for the interaction of MLO- NLuc with CLuc-CAM2 (*A. thaliana* CALMODULIN 2; positive control) on the same leaf. The *in planta* production of recombinant proteins tagged with NLuc or CLuc was assessed by immunoblotting (**Supplemental** Figure 5B). The boxplots illustrate the outcome of at least four independent experiments while individual data points represent the mean of two replicates each. Letters assign differences of statistical significance (pairwise Student’s *t*-test corrected by FDR, *α* = 0.01). **E**) Generic membrane topology of a clade V MLO protein. Protein domains that were subject to yeast two-hybrid analysis are shaded in brown. IC2, second intracellular loop; CT, carboxyl terminus. **F**) Yeast two-hybrid experiment to test the interaction of EXO70H4 with MLO6^IC2^ or MLO6^CT^. *S. cerevisiae* cultures expressing recombinant *EXO70H4* and *MLO6* constructs were dropped on a medium deprived of (1) leucine (Leu) and tryptophan (Trp) or (2) leucine, tryptophan, and histidine (His) in four different serial 1:10 dilutions. The pictures are representative of the outcome of three independent experiments (**Supplemental** Figure 6A). The production of recombinant yeast bait and prey proteins was validated by immunoblotting (**Supplemental** Figure 6B). AD, activation domain; BD, binding domain; eV, empty vector.

Based on this co-expression data, we carried out an initial screening of MLO6-EXO70 interaction by Luciferase Complementation Imaging (LCI) assays upon transient co-expression of the candidate genes in *Nicotiana benthamiana* (**Supplemental Figure 4A**). In this experiment, amino- (NLuc) and carboxyl-terminal (CLuc) fragments of firefly luciferase were fused with the proteins of interest, leading to restored luciferase activity (and consequently enhanced luminescence) upon physical interaction of the proteins under consideration (Chen et al. 2008). We used the established interaction between MLO proteins and the cytosolic calcium sensor calmodulin (Kim et al. 2002) as a positive control in this set of experiments. In the initial screening, we found a significantly enhanced (> 0.24) relative luminescence (in relation to the calmodulin positive control) for the combination of MLO6 (carboxyl-terminally tagged with NLuc) with members of the EXO70H family (amino-terminally tagged with CLuc) in comparison to most of the other EXO70 proteins tested (**Supplemental Figure 4A**). We, therefore, selected EXO70H1, EXO70H2, EXO70H4, and EXO70H7 for subsequent comprehensive analysis of protein-protein interactions including, apart from MLO6, also MLO2 and MLO12. Furthermore, we also involved EXO70B2 and EXO70D3 as controls because these two CLuc-tagged proteins showed low luminescence values (< 0.07) in the combination with MLO6-NLuc (**Supplemental Figure 4A**), despite their evident accumulation in transformed *N. benthamiana* tissue (**Supplemental Figure 4B**). In the case of MLO2-NLuc, we observed significantly enhanced relative luminescence after co-expression with CLuc-EXO70H1 (arithmetic mean ∼0.39) and -H4 (∼0.40) as opposed to CLuc-EXO70D3 (∼0.02) (**Figure 6B** and **Supplemental Figure 5A**). Strikingly, the combination of MLO2-NLuc and CLuc- EXO70B2 generated a signal with an average relative luminescence intensity of ∼0.56. This strong signal contrasts the values obtained with the MLO6-NLuc/CLuc-EXO70B2 (∼0.14) and MLO12-NLuc/CLuc-EXO70B2 (∼0.10) combinations (**Figure 6C, D**) and may hint at a preferred interaction of MLO2 with EXO70B2. In the case of MLO6-NLuc, the strongest relative luminescence was observed upon co-expression with CLuc- EXO70H4 (∼0.67) (**Figure 6C and Supplemental Figure 5A**), whereas MLO12-NLuc showed enhanced relative luminescence in combination with CLuc-EXO70H1 (∼0.37) (**Figure 6D and Supplemental Figure 5A**).

As luciferase complementation experiments indicated a direct interaction of particular EXO70 proteins with specific clade V MLO proteins, we aimed to characterize this interplay further by delimiting the interacting region within the MLO protein. For this purpose, we focused on the strong MLO6-EXO70H4 interaction (**Figure 6C**) and reasoned that the second intracellular loop (IC2) and the carboxyl terminus (CT), comprising the largest cytoplasmic regions of MLO proteins (Devoto et al. 1999) (**Figure 6E**), are the most likely parts of the protein involved in the interaction with EXO70H4. We carried out yeast two-hybrid assays and tested EXO70H4 as a bait protein against MLO6^IC2^ or MLO6^CT^ as prey proteins (**Figure 6E**). We observed the growth of yeast colonies harboring the EXO70H4/MLO6^CT^ but not the EXO70H4/MLO6^IC2^ combination on selective media lacking histidine, indicative of interaction between EXO70H4 and the MLO6 carboxyl terminus (**Figure 6F** and **Supplemental Figure 6A**).

### Combined *exo70 mlo* mutants show increased resistance to powdery mildew host cell entry

As various types of cell wall analyses and protein-protein interaction experiments indicated an interplay of EXO70H4 and MLO clade V proteins, we wondered if plants deprived of EXO70H4, alone or in combination with mutations in *MLO* genes, exhibit an altered infection phenotype upon challenge with an adapted powdery mildew pathogen (**Figure 7**). Thus, we inoculated *exo70H4* and *mlo* mutant plants as well as combined *exo70H4 mlo* mutants with conidiospores of *Erysiphe cruciferarum* and scored fungal host cell entry success on these genotypes (**Figure 7A**). Consistent with previous studies (Consonni et al., 2006; Consonni et al., 2010), we observed that about 80% of fungal sporelings successfully penetrated leaf epidermal cells of Col-0 WT plants at 72 h after inoculation (**Figure 7B**). Likewise, we scored WT-like levels of fungal host cell entry on *mlo6-2* (∼82%) and *mlo6-4* (∼84%) single mutants. Unlike *mlo6* plants, *mlo2-5* (∼63%) mutants supported significantly lower levels of host cell entry compared to Col-0 while *mlo2-5 mlo6-2 mlo12-1* triple mutants were completely resistant (∼0%) as described before (Acevedo-Garcia et al. 2017; Consonni et al. 2006). Concerning *exo70H4* single mutants, we observed lower yet not significantly reduced levels of penetration on *exo70H4-1* (∼72%) and *exo70H4-3* (∼72%) lines compared to Col-0 (∼80% - see above), pointing towards a slightly enhanced resistance phenotype of these mutants. Two different allele combinations of combined *exo70H4 mlo6* mutants, however, allowed significantly less fungal penetration (*exo70H4-1 mlo6-2* ∼ 61%, *exo70H4-1 mlo6-4* ∼62%) than Col-0 WT plants and *mlo6* single mutant lines (∼80%–84% - see above), and in tendency less host cell entry than the *exo70H4-1* single mutant line (∼72%). Similarly, *exo70H4-1 mlo2-5* double mutants were significantly less susceptible (∼ 49%) than Col-0 plants (∼80%), and the partly resistant *mlo2-5* (∼63%) and *exo70H4-1* (∼72%) single mutant lines (**Figure 7B**). Taken together, these data indicate a synergism between *exo70H4* and *mlo* mutations regarding enhanced powdery mildew resistance.

**Figure 7.**
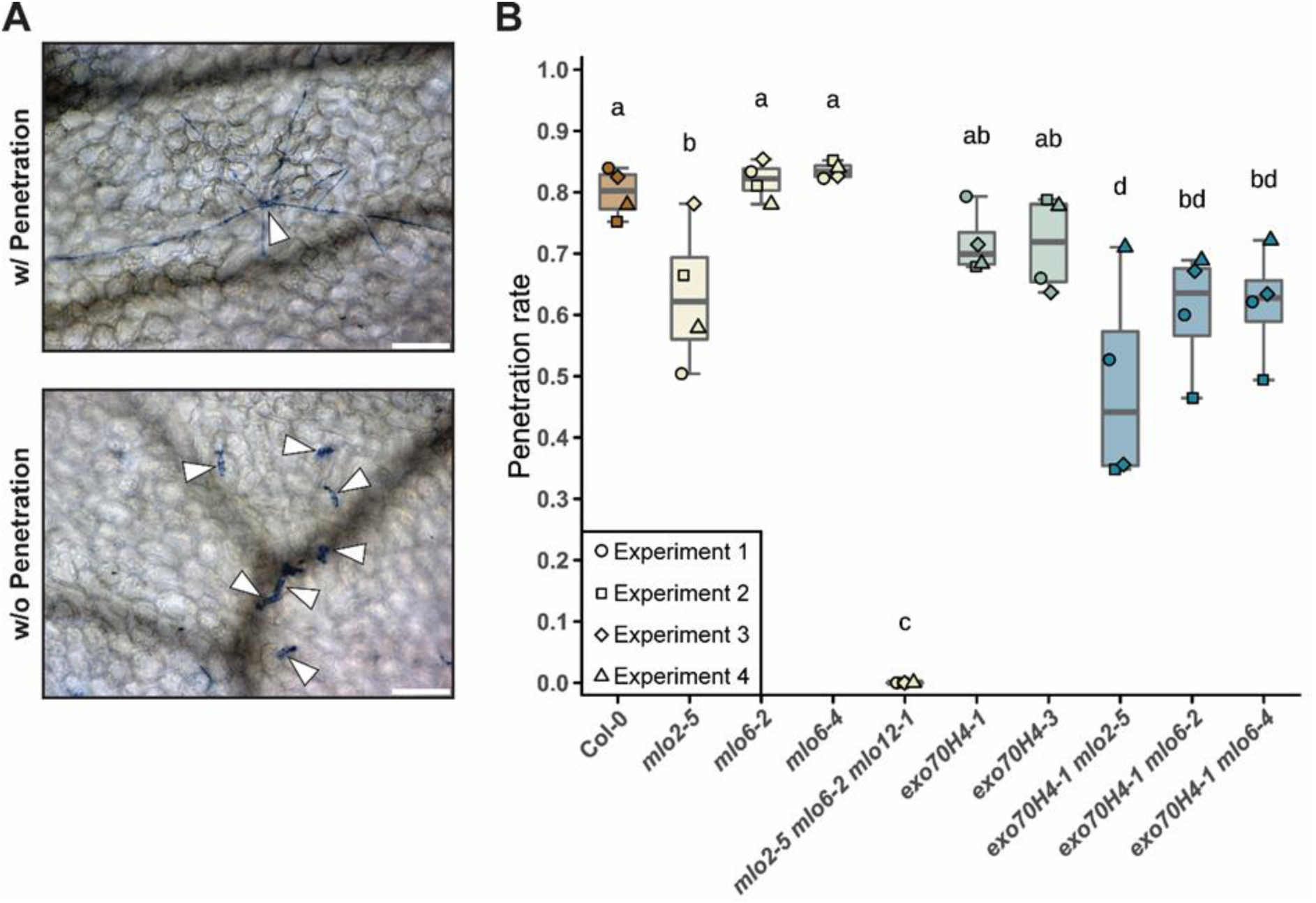
Host cell entry by *E. cruciferarum* is reduced on *exo70H4 mlo* double mutant lines. **A)** Successful and unsuccessful penetration of *E. cruciferarum* conidiospores (arrowheads) on rosette leaves of *A. thaliana*. The presence of secondary hyphae at 72 h post-inoculation reveals successful host cell penetration (w/ Penetration, top) whereas appressorium formation without the development of secondary hyphae indicates unsuccessful penetration (w/o Penetration, bottom). Fungal structures were stained with Coomassie brilliant blue. Scale bars represent 100 µm. **B)** Host cell entry rate of *E. cruciferarum* on various *A. thaliana* genotypes at 72 h post-inoculation. The boxplot illustrates the outcome of four independent experiments while data points represent the mean of three plants each. Two rosette leaves were assessed per plant, and approximately 50 fungal attack sites were scored per rosette leaf, i.e. about 1,200 sites per genotype in total. Letters assign differences of statistical significance (pairwise Student’s *t*-test corrected by FDR, *α* = 0.05).

## Discussion

We discovered a functional interplay between EXO70H4 and MLO2, MLO6, and MLO12 in the context of cell wall-related secretory processes in trichomes of *A. thaliana* rosette leaves. This claim rests on (1) a marked phenotypic overlap of *exo70H4* and *mlo2 mlo6 mlo12* triple mutants regarding secretion-dependent trichome secondary cell wall features (**Figures 1, 2 and 3**), (2) differences in trichome cell wall composition in *exo70H4* and *mlo2 mlo6 mlo12* mutant plants as revealed by biochemical analysis and/or FTIR spectroscopy of isolated trichomes (**Figure 4**), (3) extensive co-localization of fluorophore-labeled EXO70H4 and MLO2, MLO6 and MLO12 proteins (**Figure 5A**), (4) mislocalization of fluorophore-tagged EXO70H4 in *mlo2 mlo6 mlo12* triple mutant plants (**Figure 5B**) and MLO6-GFP in the *exo70H4* mutant (**Figure 5C**), (5) shared enhanced mobility of GFP-PMR4-marked intracellular compartments in transgenic *exo70H4* and *mlo2 mlo6 mlo12* mutant lines (**Figure 5D and E**; (Kulich et al., 2018)), (6) isoform-preferential interaction between EXO70 and MLO proteins (**Figure 6**), and (7) synergistically enhanced powdery mildew resistance in *exo70H4 mlo2* and *exo70H4 mlo6* double mutants (**Figure 7**). Taken together, this comprehensive dataset raises the intriguing possibility that EXO70 and MLO proteins might be involved in the same cellular pathway or even reside within the same macromolecular complex. Indeed, both the exocyst and MLO proteins have been implicated in exocytosis (Meng et al., 2020; Žárský et al., 2020; Heider and Munson, 2012), suggesting that this process is the common denominator and, therefore, likewise affected in *exo70* as well as *mlo* mutants.

While the exocyst has a well-established function in secretory processes, the role of MLO proteins therein remains enigmatic. MLO proteins were recently demonstrated to operate as PM-localized calcium channels (Gao et al., 2022). Calcium influx is a proven trigger for exocytosis in eukaryotic cells, promoting vesicle fusion events at the PM (Barclay et al., 2005). Established examples of calcium-induced secretion in mammals are synaptic vesicle exocytosis in neurons, granule exocytosis in mast cells, and hormone exocytosis in endocrine cells. Mechanistically, the calcium signal is transmitted *via* cytoplasmic calcium sensors to the membrane fusion machinery, of which synaptotagmins are arguably the best-studied ones (Pang and Südhof, 2010). In *A. thaliana*, synaptotagmin 5 was recently suggested to regulate secretory immune responses mediated by the SYP132-VAMP721/722 SNARE protein module (Kim et al., 2021). We hypothesize that PM-localized MLO calcium channels may permit calcium influx-dependent and possibly spatially confined secretory events, which also involve the physical association of MLO proteins with EXO70 proteins. The protein-protein interaction between a member of the exocyst complex and MLO calcium channels might determine the actual site of vesicle fusion events, thereby allowing for fine-tuned polar exocytosis. This scenario is compatible with delocalized exocytosis in *exo70* and *mlo* mutant plants, resulting in stochastic patterns of secondary cell wall biogenesis in trichomes, as observed in our experiments (**Figure 1-3**). The shared phenotype of *exo70H4* and *mlo* triple mutants reinforces the notion that the interaction between EXO70 and MLO proteins (**Figure 6**) is biologically relevant.

If this scenario was true, one may wonder what could be the cargo of the secretory pathway linked to an EXO70H4-MLO regulatory module in trichomes. A prime candidate is the PMR4 callose synthase, which along with CalS9 was previously shown to be mislocalized in the *exo70H4-1* mutant (Kulich et al., 2018) and for which we likewise observed a similar mislocalization pattern in the *mlo2-5 mlo6-2 mlo12-1* triple mutant (**Figure 5D and E**). This mislocalization of PMR4 is characterized by a reduction of immobile PM-associated GFP-PMR4 speckles and a concomitant increase in mobile GFP-PMR4-labelled membrane bodies, pointing to a reduction of PMR4 secretion in *exo70* and *mlo* triple mutants. Additional cargos of the proposed exocyst-MLO secretory pathway that would account for the autofluorescence (possibly phenolic compounds; (Kulich et al., 2018)), ROS accumulation and heavy metal deposition (reported previously to depend on EXO70H4; (Kulich et al., 2018)) in trichomes are still unknown. One can, however, speculate about their nature. The zinc ZIP transporters are highly expressed in trichomes (Jakoby et al., 2008), and a copper transporter (COPT5, At5g20650) was found to be the most enriched protein in a comprehensive trichome proteome analysis (Huebbers et al., 2022). Recent studies reported that zinc accumulates at the basal part of the trichome (Kulich et al., 2015; Ricachenevsky et al., 2021), roughly corresponding to the region of the OR. Therefore, zinc and other heavy metal transporters can be considered additional candidates for the EXO70H4-MLO secretory pathway. *A. thaliana* trichomes contain two distinct PM domains, with EXO70H4 and specific phospholipids being enriched in the apical one (Kubátová et al., 2019). We, thus, believe that the basal localization of zinc in trichomes as described before (Ricachenevsky et al., 2021; Kulich et al., 2015) represents part of the basal domain defined by the absence of EXO70H4 and the presence of EXO70A1 (Kulich et al., 2018; Kubátová et al., 2019). Our HM staining experiments as well as EXO70H4 localization data further support this notion.

Interestingly, while EXO70H4 is involved in all the aspects of trichome cell wall maturation considered in the present study, the contribution of the three MLO proteins seems to vary. Previous studies already indicated isoform-specificity for *mlo*- conditioned mutant phenotypes. For example, *A. thaliana* genes *MLO2*, *MLO6,* and *MLO12* confer susceptibility to powdery mildew fungi (Consonni et al., 2006). In the context of this phenotype, the three genes exhibit unequal redundancy, with *MLO2* being the major player, resulting in vastly different outcomes for *mlo2* single mutants (partial resistance) and *mlo2 mlo6 mlo12* triple mutants (full resistance; (Consonni et al., 2006). Similarly, enhanced tolerance to ozone is specifically conditioned by loss-of-function *mlo2* mutants and further increased in a *mlo2 mlo6 mlo12* triple mutant (Cui et al., 2018). By contrast, both *mlo2* single mutants and *mlo2 mlo6 mlo12* triple mutants show a complete loss of systemic acquired resistance, albeit *MLO6* may have a supportive but less critical role in this type of plant defense (Gruner et al., 2018). Based on respective mutant phenotypes, we here found a major contribution of *MLO6* regarding trichome callose deposition (**Supplemental Figure 1A**) and cell wall thickness (**Supplemental Figure 1B**) as well as a possible joint function for *MLO2* and *MLO6* regarding trichome cell wall autofluorescence (**Supplemental Figure 1D**) and HM deposition (**Supplemental Figure 1E**). The predominant role of *MLO6* in trichome-associated processes is consistent with the high level of expression of this gene in trichomes (Jakoby et al., 2008) and the favored interaction of MLO6 with the likewise preferentially in trichomes expressed EXO70H4 protein (**Figure 6A**; (Jakoby et al., 2008)). However, at least for some of the trichome phenotypes, e.g. callose deposition (**Supplemental Figure 1A**) and ROS accumulation (**Supplemental Figure 1C**), it seems as if the severity of the phenotype in the *mlo* triple mutants exceeds that of the *mlo6* single and/or *mlo2 mlo6* double mutants, suggesting that also MLO12 contributes to these features. Thus, although *MLO6* appears to be the main player, all three *MLO* genes may contribute to different degrees and with varying redundancy to the various trichome phenotypes.

Regarding some of the trichome phenotypes, we noticed a weaker penetrance for the *mlo2-6 mlo6-4 mlo12-8* triple mutant as compared to the *mlo2-5 mlo6-2 mlo12-1* triple mutant. This applies to cell wall thickness (**Figure 1E**), ROS accumulation (**Figure 2B**), and HM deposition (**Figure 2D**) in leaf-attached trichomes as well as the histochemical staining patterns for calcofluor white (**Figure 3A**), ruthenium red (**Figure 3B**) and toluidine blue O (**Figure 3C**) in detached trichomes. It could be further noticed regarding global cell wall composition as assessed by FTIR spectroscopy analysis (**Figure 4D**) and the levels of the monosaccharides arabinose (**Figure 4A**) and galactose (**Figure 4B**). A weaker phenotype has been already previously described for the *mlo2-6 mlo6-4 mlo12-8* triple mutant regarding powdery mildew resistance, exemplified by some residual susceptibility to the fungal pathogen (Acevedo-Garcia et al., 2017), which differs from the complete resistance of the *mlo2-5 mlo6-2 mlo12-1* triple mutant (Consonni et al., 2006). The differential outcome of the two *mlo* triple mutants in powdery mildew pathoassays has been ascribed to the rather distal insertion site of the T-DNA in the *mlo6-4* allele, possibly resulting in some residual MLO6 function in the context of this mutant and, accordingly, also in the *mlo2-6 mlo6- 4 mlo12-8* triple mutant (Acevedo-Garcia et al., 2017). It is, thus, tempting to speculate that trichome phenotypes with a differential manifestation for the two *mlo* triple mutants are mainly caused by a loss of function of *MLO6*.

Our studies demonstrated physical interactions between EXO70 and MLO proteins in two different *in vivo* experimental systems: *in planta* LCI assays and yeast two-hybrid assays (**Figure 6B–D, Figure 6F, Supplemental Figure 5,** and **Supplemental Figure 6**). The LCI experiments revealed preferential interaction of particular EXO70 and MLO isoforms, yet often also indicated weaker interactions with the non-preferred paralogs. This outcome may point towards some level of promiscuity regarding EXO70-MLO interactions, which may at least in part explain the genetic redundancies observed for some *exo70* and *mlo* mutant phenotypes. The yeast two-hybrid assay delimited the interacting region of MLO6 to its carboxyl-terminal cytoplasmic domain (**Figure 6F**), which also harbors the conserved calmodulin-binding domain (Kim et al., 2002; Kusch et al., 2016). This section of MLO proteins was recently shown to serve as a regulatory center, modifying the calcium channel activity of MLO proteins in a calcium-calmodulin-dependent autoinhibitory manner (Gao et al., 2022). It will be interesting to study in the future whether the binding of calmodulin to MLO proteins also affects its interaction with EXO70 proteins. It will be likewise informative to dissect EXO70 proteins to find out which parts of these polypeptides associate with MLO proteins.

Our pathoassays revealed a synergistic contribution of *exo70H4* and *mlo2* mutants regarding powdery mildew resistance. While single mutants of these two genes showed a moderate decrease in susceptibility (host cell entry) to the fungal pathogen, an *exo70H4 mlo2* double mutant exhibited an additive reduction in host cell penetration (**Figure 7B**). Notably, we observed a similar synergism for *exo70H4 mlo6* double mutants (**Figure 7B**). The latter outcome is even more surprising, as *mlo6* single mutants do not exhibit any reduction in powdery mildew host cell entry rates (**Figure 7B;** (Consonni et al., 2006)). However, in combination with *mlo2* mutants, *mlo6* (and also *mlo12*) mutations enhance the *mlo2* pathogen phenotype, indicating unequal genetic redundancy of *MLO2*, *MLO6,* and *MLO12* regarding the modulation of powdery mildew susceptibility (Consonni et al., 2006). Reminiscent of this scenario, a similar cryptic contribution of the *mlo6* mutation becomes apparent in combination with the *exo70H4-1* mutant (**Figure 7B**). In summary, this data uncovers a critical role for EXO70H4 in plant defense against an adapted powdery mildew pathogen. We hypothesize that similar to MLO2, MLO6, and MLO12, further EXO70 paralogs possibly contribute redundantly to this process. This notion is supported by the expression profiles of *EXO70* genes upon biotic stresses (**Figure 6A**), which unraveled further *EXO70* paralogs as highly expressed under these conditions. Higher order *exo70* mutants, with or without additional *mlo* mutations, will be required to disentangle the putative overlapping and/or redundant functions of EXO70 proteins in plant defense against powdery mildew fungi. We hypothesize that EXO70 and MLO proteins jointly regulate secretory processes in the context of plant immunity, including powdery mildew challenge. Cellular polarization and polar secretion have long been suggested to be decisive in antifungal defense (Schmelzer, 2002; Kwon et al., 2008b; Yun et al., 2008). Genetic studies unraveled indeed components of the secretory machinery such as SNARE proteins and ABC transporters as being crucial for effective preinvasive immunity against adapted and non-adapted powdery mildew fungi, also in the context of *mlo*-based resistance (Kwon et al., 2008a; Collins et al., 2003; Bednarek et al., 2009; Consonni et al., 2006). The herein described identification of EXO70 subunits as additional factors involved in this process further highlights the presumed role of polar secretion in antifungal defense at the cell wall. Deregulated secretory processes in *mlo* and *exo70* mutants may result in a constitutively altered cell wall architecture in these mutants, which might in part account for their altered pathogen infection phenotypes. Recently reported direct interactions of both MLOs and plant exocyst subunits (including EXO70s) with SNARE proteins reinforce the conclusion that both types of proteins are part of a hierarchical module regulating exocytosis (Meng et al., 2020; Ortmannová et al., 2022; Larson et al., 2020). Although we consider the PMR4 callose synthase as an important cargo of EXO70-MLO cooperation in the context of trichome secondary cell walls, PMR4 is unlikely to play a major role regarding powdery mildew resistance as a previous study demonstrated that *mlo2*-mediated resistance is independent of PMR4-based callose deposition (Consonni et al., 2010).

An involvement of EXO70 proteins in plant immunity has been described before. For example, *A. thaliana* EXO70B1 and EXO70B2 were previously reported by several studies to be linked to pathogen defense. *exo70b1* mutant plants exhibited altered infection outcomes upon challenge with various pathogens, associated with a lesion-mimic cell death phenotype and reduced responsiveness to microbe-derived molecules (Stegmann et al., 2013; Zhao et al., 2015). The activated defense responses of the *exo70B1* mutant rely on a truncated resistance protein of the NLR class (TN2), which might monitor the integrity of EXO70B1 (Zhao et al., 2015). Furthermore, *exo70B2* and *exo70H1* mutants showed increased susceptibility to the bacterial pathogen *Pseudomonas syringae* pv. *maculicola*. The *exo70B2* mutant exhibited, in addition, the formation of aberrant local cell wall appositions (papillae) in response to inoculation with the barley powdery mildew pathogen, *Blumeria hordei* (Pecenkova et al., 2011; Ortmannová et al., 2022). Finally, both EXO70B1 and EXO70B2 regulate the trafficking of the pattern recognition receptor FLS2, which recognizes bacterial flagellin, to the PM (Wang et al., 2020). Apart from *A. thaliana*, EXO70 proteins were linked to plant immunity in several other plant species. In barley, for example, a member of a lineage-specific EXO70 clade, EXO70FX12, is required for receptor kinase-mediated disease resistance to the wheat stripe rust pathogen, *Puccinia striiformis* f.sp. *tritici* (Holden et al., 2022). It is, thus, not surprising that defense-related EXO70 subunits such as EXO70B1 are targets of pathogen effector molecules that affect their function (Michalopoulou et al., 2022; Wang et al., 2019). It will be interesting to study whether effector proteins of powdery mildew fungi target EXO70 subunits and/or MLO proteins for promoting virulence.

Though we provide in our work strong evidence for an EXO70-MLO interplay, we appreciate that our current data do not provide any evidence for a contribution of the exocyst core subunits, although we previously showed in a yeast two-hybrid assay that EXO70H4 does interact with the *A. thaliana* exocyst core subunits SEC5a, SEC6 and EXO84b (Kulich et al. 2015). It, therefore, remains a formal possibility that the newly discovered EXO70-MLO interaction relates to cellular functions other than secretory processes, involving non-canonical EXO70 activities without the whole exocyst complex. However, we consider this scenario rather unlikely given that so far plant EXO70 subunits were exclusively linked to functions in exocytosis and because both *exo70H4* and *mlo* mutants exhibit trichome mutant phenotypes that are indicative of perturbed secretion. Further work will be required to unravel whether the herein proposed EXO70-MLO module also operates in other biological processes in which EXO70 and MLO proteins have been implicated before.

## Materials and Methods

### Plant material

The *A. thaliana exo70H4-1* (SALK_023593; (Kulich et al., 2015) mutant and the *mlo2- 5 mlo6-2 mlo12-1* (Consonni et al., 2006) and *mlo2-6 mlo6-4 mlo12-8* (Acevedo-Garcia et al., 2017) triple mutants, as well as the respective *mlo* single and double mutants, are T-DNA/transposon insertion mutants in the background of ecotype Col- 0.

### Plant cultivation

*A. thaliana* and *N. benthamiana* plants were sown on SoMi 531 soil (HAWITA, Vechta, Germany). Seeding and harvest of *A. thaliana* that were subjected to trichome isolation were carried out as described before (Huebbers et al., 2022). For all experiments, plants were cultivated at 22/20 °C average day/night temperature with a photoperiod of 10 h d^−1^, a photosynthetic photon flux density of 80–100 μmol m^−1^ s^−1^, and relative humidity of 80–90%. Before leaf-to-leaf touch inoculation with *E. cruciferarum*, 35-d old *A. thaliana* plants were placed in a separate growth chamber with a constant temperature of 20 °C and relative humidity of 68%. Photoperiod and photosynthetic photon flux density were the same as described before. Leaves of inoculated *A. thaliana* plants were sampled at 72 h post-inoculation. After infiltration of *Agrobacterium tumefaciens*, *N. benthamiana* plants were transferred to a photoperiod of 16 h d^−1^ with a photosynthetic photon flux density of 105–120 μmol m^−1^ s^−1^ as well as an average day/night temperature of 23/20 °C and relative humidity of 80–90%. Post-infiltration incubation was carried out for three days. For the analysis of callose, autofluorescence, HM, and ROS phenotypes, the plants were cultivated on Jiffy pellets (Jiffy Products International BV, Zwijndrecht, The Netherlands) at 23/20 °C average day/night temperature with a photoperiod of 16 h light and 8 h dark.

### Histochemical staining of leaf-attached trichomes for callose staining and autofluorescence

The staining of callose in *A.* trichomes was performed as described previously (Kulich et al., 2018). For the observation of autofluorescence, the third or fourth visible leaf was used. The leaf was squeezed on the microscope slide and the trichomes were brushed with the help of a needle; then observed after 5-10 min without a cover glass.

### Histochemical staining of leaf-attached trichomes for ROS

The detection of hydrogen peroxide was conducted as reported before (Daudi and O’Brien, 2012). Third or fourth leaves of three-week-old plants were used. The staining was performed one hour after cutting, to stabilize the ROS level in the whole leaf. Leaves were vacuum-infiltrated for 5 min with the DAB solution and then incubated on a shaker for 5 h. Following the incubation, the leaves were transferred to 50 ml test tubes with a bleaching solution (3:1:1 mixture of ethanol, acetic acid, and glycerol) and boiled for 15-20 min (until leaves were completely bleached). Following bleaching, leaves were placed in 6-well plates and observed the day after.

### Histochemical staining of leaf-attached trichomes for HMs

Three-week-old *A. thaliana* plants watered with 1 mM of zinc chloride solution for one week were stained with dithizone as described previously (Seregin and Ivanov, 1997). The staining solution was composed of 30 mg of dithizone dissolved in 20 ml of distilled water, 60 ml of acetone, and 8 drops of acetic acid. The leaves were stained for one hour and observed the same day. Before observation, each leaf was rinsed in distilled water.

### Trichome isolation

Trichome release and enrichment were carried out according to the STIRRER method introduced previously (Huebbers et al., 2022). Briefly, freshly harvested *A. thaliana* seedlings were transferred to a 500-mL beaker containing 250 mL of phosphate-buffered saline (PBS; 10 mM disodium hydrogen phosphate, 2.7 mM potassium chloride, 2.0 mM potassium dihydrogen phosphate, 0.5 mM magnesium chloride, 137 mM sodium chloride, pH 7.5) solution supplemented with 50 mM ethylenediaminetetraacetic acid (EDTA). The trichome buffer suspension was stirred at 300 rpm for 30 min. Subsequently, four layers of screen door mesh were used to separate the processed seedlings and the buffer solution containing the released trichomes. The latter was captured in a 300 mL beaker and poured through a cell strainer (VWR, Radnor, USA) with a nominal pore size of 100 µm. The restrained trichomes were collected by a small spatula and transferred to 2-mL reaction tubes. After the first round of agitation and filtration, the process was repeated two times for 15 min, each. Enriched trichomes were either stored at 4 °C in 1 mL phosphate-buffered saline buffer or frozen in liquid nitrogen and lyophilized for monosaccharide and cellulose quantification or FTIR spectroscopy.

### Histochemical staining of isolated trichomes

Histochemical staining of *A. thaliana* trichomes was carried out using 20 µL of the respective trichome suspension sample, which was mixed to equal shares with the desired staining solution. Incubation took place in the dark. For calcofluor white, trichomes were incubated for 60 min and samples were observed by UV illumination. For ruthenium red and toluidine blue O, samples were incubated for 120 min and subjected to bright-field illumination. Calcofluor white, ruthenium red, and toluidine blue O solutions were prepared as described previously (Pradhan Mitra and Loqué, 2014).

### Quantification of neutral sugars and cellulose

The mass of neutral monosaccharides and cellulose in isolated and lyophilized trichomes was determined as described (Yeats et al., 2016). In brief, alcohol-insoluble material was prepared from the dried trichomes and split into two samples – one sample was treated with a weak acid (4% sulfuric acid) to release matrix polysaccharide-derived sugars, while the other sample was treated with a strong acid (72% sulfuric acid) to swell cellulose followed by the weak acid (4% sulfuric acid) to yield monosaccharides derived from cellulose and the matrix polymers. Subtraction of the two values allows for the quantification of crystalline cellulose. Monosaccharides were quantified by high-performance anion-exchange chromatography with a pulsed amperometric detector (HPAEC-PAD).

### Quantification of uronic acids

Isolated and lyophilized trichomes were acid hydrolyzed with 2-N-trifluoroacetic acid at 121 °C and dried under nitrogen flow at 30 °C. Then the samples were resuspended in miliQ water and assessed by the m-hydroxybiphenyl method (Blumenkrantz and Asboe-Hansen, 1973) using galacturonic acid as a standard.

### FTIR spectroscopy

FTIR spectra were obtained from isolated and lyophilized trichomes using a JASCO4700 FTIR spectrometer (JASCO, Pfungstadt, Germany) equipped with an attenuated total reflection module (ATR) at a resolution of 1 cm^-1^. The region between 800 and 1800 cm^-1^ was selected from the average spectra (n = 10) to analyze differences in cell wall components (Alonso-Simón et al., 2011). Subsequently, all spectra were normalized and baseline-corrected by using Spectra Manager (JASCO).

### Generation of plant expression constructs and transgenic lines

The mCherry-EXO70H4 construct under the control of the native *EXO70H4* promoter and GFP-PMR4 expressed from the ubiquitin promoter were described before (Kulich et al., 2015; Kulich et al., 2018). *MLO2* and *MLO6* cDNA and *MLO12* genomic sequences were amplified by PCR using suitable primers and integrated *via* BP recombination into Gateway^®^-compatible pDONR221 entry vectors (Thermo Fisher Scientific, Darmstadt, Germany). Inserts and the *EXO70H4* were co-shuttled *via* multi- site Gateway^®^ recombination into plant expression vectors pK7m34GW ((Karimi et al., 2005); *MLO2*) or pH7m34GW ((Karimi et al., 2005); *MLO6* and MLO12). Similarly, an MLO6-GFP plant expression construct under the control of the *MLO6* promoter was established by multi-site Gateway^®^ recombination. The resulting binary expression vectors were transferred into *Agrobacterium tumefaciens* strain GV3101 (pMP90RK) and transgenic *A. thaliana* lines generated *via* the floral dip method (Clough and Bent, 1998). The selection of transgenic lines was performed on suitable selection media.

### Microscopy

The observation of callose and autofluorescence for leaf-attached trichomes was done on a Nikon Eclipse 90i microscope with either a PlanApo 4×/0.2 objective (for the quantification of callose and the acquisition of autofluorescence) or a PlanApo 10x/0.45 objective (for imaging single trichomes). Micrographs were captured with a Nikon DsFi 2 camera. For the observation of ROS, an Olympus IX71 microscope with a LUCPlanFLN 40x/0.6 objective was used. For imaging HM deposition, a Nikon Eclipse 90i microscope with a PlanApo 20x/0.75 objective was used. Trichome cell wall thickness was observed with an Olympus BX51 microscope with a UPlanSApo 60x/1.20 objective with water immersion. Micrographs of histochemically stained isolated trichomes were captured using a BZ-9000 transmitted light and epifluorescence microscope (Keyence, Osaka, Japan).

For confocal microscopy, a Zeiss LSM880 with a C-Apochromat 40/1.2 W correction FCS M27 objective and a Leica SP8 with a 60x water correction objective were used. Excitation (in brackets) and emission wavelengths were as follows: GFP (488 nm) 508–540 nm, chlorophyll (488 nm) 650–721 nm, cell wall autofluorescence (405 nm) 426–502 nm, mCherry (561 nm) 597–641, as described before (Kulich et al., 2018).

### Image analysis

All micrographs of leaf-attached trichomes were processed using the Fiji platform. To calculate the number of stable PM-associated GFP-PMR4-marked “dots” (compartments) in *A. thaliana* trichomes, we imaged the trichomes in a time series experiment and subsequently performed the minimal projections command in the Fiji software (Schindelin et al., 2012; Kulich et al., 2015). We then calculated the number of visible “dots” using the dot and ROI Manager tools. Image optimization for micrographs of histochemically stained isolated trichomes was carried out identically for all micrographs of the same staining type using Adobe Photoshop 2022.

### Luciferase complementation imaging

*N. benthamiana* plants were used for transient gene expression with *A. tumefaciens*, harboring the appropriate constructs and the p19 silencing suppressor plasmid, as described before (Campe et al., 2016) with minor modifications. *A. tumefaciens* strain GV3101 (pMP90RK) was transformed with plasmids harboring different *EXO70* and *MLO* coding sequences that were transferred from pDONR207 entry clones into pAMPAT-CLuc-GWY and pAMPAT-GWY-NLuc vectors (Gruner et al., 2021) *via* Gateway^®^ recombination. For leaf infiltration of agrobacteria, bacterial cultures were grown overnight and resuspended in an infiltration medium (10 mM MES, pH 5.6, 10 mM MgCl_2_, 150 µM acetosyringone), adjusted to an OD_600_ of 0.5, and incubated at room temperature for 2 h. For co-infiltration, equal volumes of each *A. tumefaciens* strain were mixed and infiltrated into the abaxial side of fully expanded leaves of four to six-week-old *N. benthamiana* plants with a syringe lacking a cannula. Split-luciferase complementation was assessed at 2 days after infiltration by spraying the transformed leaves of *N. benthamiana* with 1 mM D-luciferin (PerkinElmer, Rodgau, Germany) solved in water supplemented with 0.01% (v/v) Tween-20. Leaves were kept in the dark for 10 min before luminescence was detected with a ChemiDoc XRS+ imaging system (BioRad, Feldkirchen, Germany). Luminescence intensities mm^-2^ were evaluated using the Image Lab software (BioRad, Feldkirchen, Germany).

### Yeast two-hybrid assays

For yeast-two-hybrid assays, *S. cerevisiae* was transformed with plasmids expressing EXO70H4 and either the IC2 or CT fragment of MLO6, respectively. The MLO6 IC2 fragment (MLO6^IC2^) comprises MLO6 amino acids 183 to 284, and the MLO6 CT fragment (MLO6^CT^) covers MLO6 amino acids 433–583. For cloning of these constructs, *EXO70H4*, *AtMLO6^IC2^*, and *AtMLO6^CT^* coding sequences were mobilized from pDONR207 entry clones via Gateway^®^ recombination into yeast two-hybrid vectors pGADT7 and pGBKT7. The recombinant plasmids were transformed by the LiAc/PEG method (Gietz and Woods, 2002) into *S. cerevisiae* strain AH109. Yeast transformants expressing the recombinant proteins (see below) were dropped on a medium lacking leucine and tryptophan for growth control, and on a medium lacking leucine, tryptophan, and histidine to detect putative interactions. For the drop tests, liquid yeast cultures were grown overnight in a liquid medium at 28 °C and 250 rpm. The cells were harvested the following day by 1 min of centrifugation at 3,000 *g* and washed with sterile water before adjusting the OD_600_ of the solutions to 1. Ten-fold dilution series were established over four orders of magnitude and 4 µL per dilution and construct combination were dropped onto the aforementioned media.

### Total phenolic protein extraction

Frozen *N. benthamiana* leaf tissue samples were homogenized to carry out a total phenolic protein extraction. The phenolic total protein extraction was performed as described before (Thomas et al., 2015) with minor changes. The homogenized tissue samples were washed twice with ice-cold acetone and the samples were pelleted between washing steps at 16,000 *g* for 5 min at 4 °C, before being resuspended in 10% (m/v) trichloroacetic acid (TCA) in acetone. After resuspension, the samples were transferred into an ultrasonic ice water bath for 10 min. Following sonification, the samples were pelleted again for 5 min at 16,000 *g* (4 °C). The pellets were washed once with 10% (m/v) TCA in acetone, once with 10% (m/v) TCA, and then with 80% (v/v) acetone, samples were resuspended and pelleted (16000 *g*, 4 °C, 5 min) between washing steps. Shortly air-dried pellets were resuspended in freshly prepared dense sodium dodecyl sulfate (SDS) buffer (100 mM Tris-HCl, pH 8.0; 30% (m/v) saccharose; 2% (m/v) SDS; 50 mM dithiothreitol) at room temperature, before adding Tris-phenol and thoroughly mixing the samples until they become whitish. The upper phenolic phase was separated by centrifugation (16,000 *g*, 20 min, room temperature) and proteins were recovered by precipitation for 60 min at -20 °C with 5 volumes of 100 mM (v/v) ammonium acetate in methanol. After a final centrifugation step (16,000 *g*, 5 min, 4 °C), the protein pellets were washed once with 100 mM (v/v) ammonium acetate in methanol and once with 80% (v/v) acetone.

### SDS-PAGE and immunoblot analysis

The separation of the proteins was carried out *via* sodium dodecyl sulfate-polyacrylamide gel electrophoresis (SDS-PAGE). Protein samples were resuspended in NuPAGE LDS Sample buffer (Thermo Fisher Scientific) and denatured by boiling at 95 °C for 5 min before gel loading. Subsequently, 5 µg of total protein was subjected to SDS-PAGE, transferred to a nitrocellulose membrane, and used for immunoblot detection according to the manufacturer’s instructions. For luciferase fusion proteins, an α-luciferase antibody (Merck KGaA, Darmstadt, Germany) was used in a 1000^-1^ dilution in 5% (m/v) milk in Tris-buffered saline with Tween-20 (TBST; 0.1% Tween- 20, 0.14 M sodium chloride, 0.02 M Tris, pH 7.6). For Gal4 fusion proteins, α-Gal4BD and α-Gal4AD antibodies (both Santa Cruz Biotechnology, Dallas, Texas, USA) were used in a 1,000^-1^ dilution in 5% (m/v) milk in TBST. A goat or mouse α-rabbit antibody coupled to horseradish peroxidase (Santa Cruz Biotechnology Inc., Dallas, USA) in a 2000^-1^ dilution in 5% (m/v) milk in TBST was used as the secondary antibody to detect luciferase and Gal4 fusion proteins, respectively. Chemiluminescence detection of antigen-antibody complexes was carried out with SuperSignal West Femto Western substrate (Thermo Fisher Scientific). As a loading control, the nitrocellulose membrane was stained with Ponceau S (AppliChem GmbH, Darmstadt, Germany) solution.

### Computation and statistical procedures

R v.4.1.3 (R foundation, www.r-project.org/) was used for plotting, statistical analyses, and filtering of data during this study. Plotting was carried out using the ggplot2 library whereas data processing was facilitated by the dplyr library both included in the Tidyverse package (Wickham et al., 2019). Boxplots were assembled in the style of Tukey: The boundaries of the whiskers are defined by the lowest or highest value, respectively, within a 1.5 times interquartile range. The box marks the range between the first and third quartile. Horizontal bars represent the median. Note that in the text the arithmetic mean was used to compare different samples. Analysis of FTIR spectra by PCA was accomplished using factoextra (Kassambara and Mundt, 2020) and FactoMineR (Lê et al., 2008). Absolute (tissue-specific) or relative (biotic stress) transcript data were retrieved from http://bar.utoronto.ca/efp/cgi-bin/efpWeb.cgi on 2021-04-19. Correlation analysis of these transcript data was facilitated by the corrplot package (Wei and Simko, 2021). We assessed the distribution of our data by quantile-quantile plots and by the Shapiro-Wilk test (*α* = 0.05; (Shapiro and Wilk, 1965)) and checked for homoscedasticity by Levene’s test (*α* = 0.05; (Gastwirth et al., 2009)). Normally distributed data with similar variances were subjected to Student’s *t*-test (Student, 1908), whereas statistical differences between non-normally distributed samples were reviewed by the Wilcoxon-Mann-Whitney test (Mann and Whitney, 1947). Pearson’s chi-squared test (Pearson, 1900) was used to assess the statistical differences between nominal data. In the case of multiple comparison hypothesis testing, FDR was used to correct for the increased probability of type I errors (Benjamini and Hochberg, 1995).

## Supplemental Figures

Supplemental Figure 1. Quantification of trichome phenotypes in *exo70H4-1* and *mlo* mutants.

Supplemental Figure 2. *A. thaliana exo70H4* and *mlo2 mlo6 mlo12* triple mutants have altered trichome cell wall characteristics.

Supplemental Figure 3. Co-localization details of mCherry-EXO70H4 and MLO12- GFP in trichomes of transgenic lines.

Supplemental Figure 4. Analysis of the interaction of MLO6 with various EXO70 proteins by LCI.

Supplemental Figure 5. Representative luminescence signals and validation of protein production in leaves of *N. benthamiana* after luciferase complementation imaging.

Supplemental Figure 6. Yeast two-hybrid assays suggest an interaction of EXO70H4 with the carboxyl terminus MLO6.

## Acknowledgments

We thank Franz Leissing for suggestions and critical discussions regarding the project.

## Author contributions

JWH carried out histochemical staining of isolated trichomes, prepared trichome samples for monosaccharide and cellulose quantification and FTIR spectroscopy, analyzed cell wall and transcript data, performed luciferase complementation imaging and yeast-two-hybrid experiments, carried out immunoblot analysis, assessed the powdery mildew infection phenotypes of *A. thaliana* WT and mutant plants, designed the Figures and performed statistical analysis GAC generated constructs; established transgenic lines, accomplished histochemical staining of leaf-attached trichomes, performed microscopy, and conducted image analysis and statistical analysis ZV participated in the planning of the experiments, cloned constructs for the visualization of fluorophore-labeled proteins, generated and characterized transgenic lines, performed microscopy, characterized phenotypic deviations in mutant trichomes, assisted in data interpretation

PS generated transgenic lines, performed microscopy, and conducted image analysis

SCJL performed luciferase complementation imaging assays and conducted the related immunoblot analysis

HK discovered the phenotypic overlap between *mlo2 mlo6 mlo12* triple mutants and the *exo70H4* mutant

IK contributed to study conception together with HK

AR generated constructs and selected higher order mutant lines

KB contributed to trichome sampling for biochemical carbohydrate measurements and established histochemical staining experiments with isolated trichomes

AMR and HMM performed

FTIR experiments and analyzed uronic acid content

MP performed biochemical analysis of neutral carbohydrates and cellulose

RP and VZ conceived the study

GAC, PS, JWH, and RP drafted the manuscript

## Data availability statement

All relevant data can be found within the manuscript and its supporting materials.

## Abbreviations

CaM: Calmodulin
DAB: 3,3’-diaminobenzidine
EXO: Exocyst subunit
FDR: False discovery rate
FTIR: Fourier transform infrared
HM: Heavy metal
LCI: Luciferase complementation imaging
MLO: Mildew resistance locus o
OR: Ortmannian ring
PCA: Principle component analysis
PM: Plasma membrane
ROS: Reactive oxygen species
SDS: Sodium dodecyl sulfate
WT: Wild type

## Supplemental Figures

**Supplemental Figure 1.**
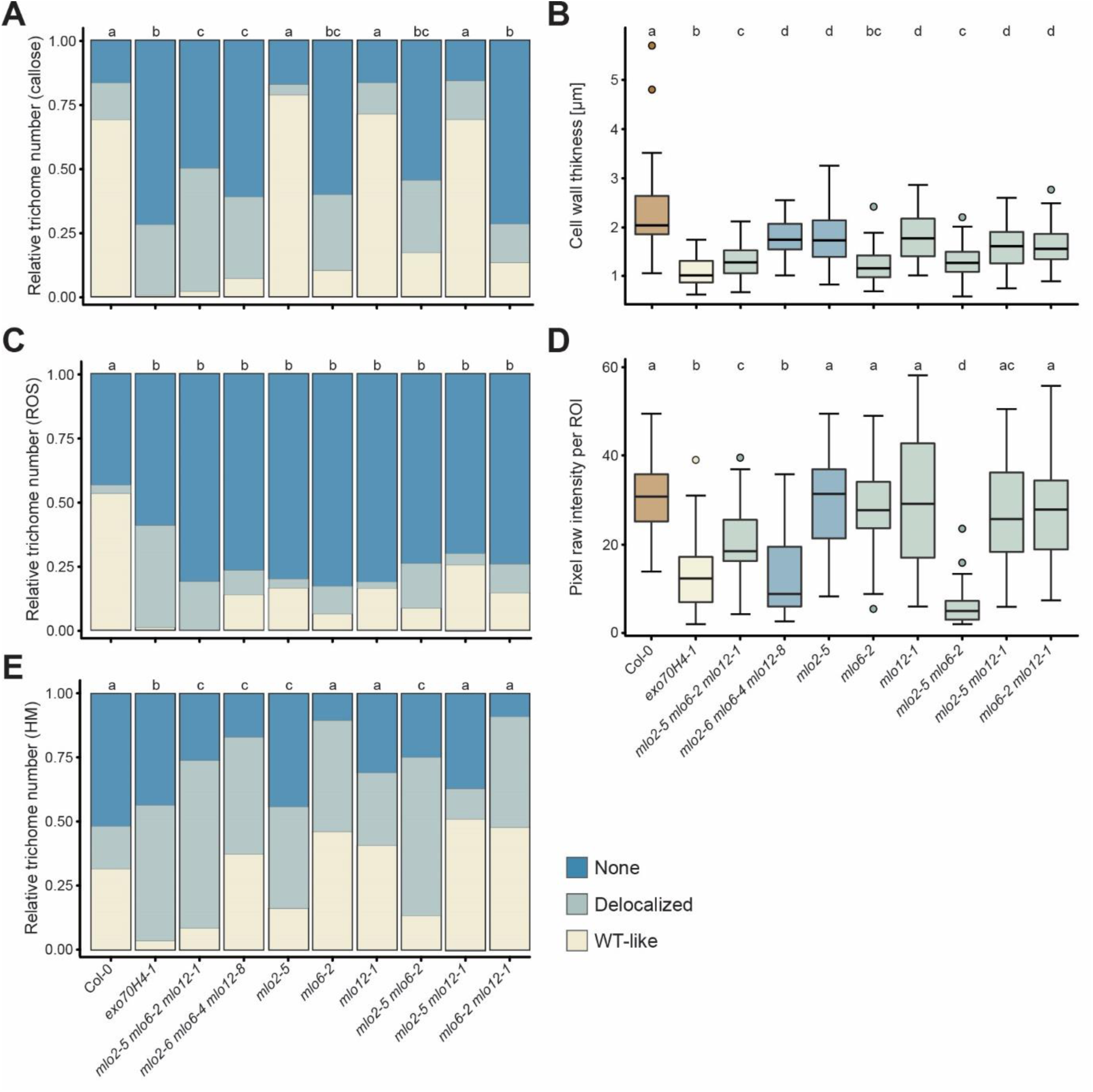
Quantification of trichome phenotypes in *exo70H4-1* and *mlo* mutants. Quantification of callose deposition (**A**), cell wall thickness (**B**), ROS accumulation (**C**), cell wall autofluorescence (**D**), and HM deposition (**E**) in trichomes of Col-0 WT, *exo70H4-1* and various *mlo* single, double and triple mutants. Values are based on three experiments with approximately 50 trichomes inspected per experiment and genotype. Data in **A**, **C**, and **E** were analyzed by chi-square test, and the *p*-values were corrected by FDR (*α* = 0.05). Data in **B** and **D** were analyzed by pairwise Wilcoxon-Mann-Whitney test and *p*-values were corrected by FDR (*α* = 0.01). Letters assign differences of statistical significance. Note that the values shown for Col-0 WT, the *exo70H4-1* single mutant, and the two *mlo2 mlo6 mlo12* triple mutants are identical to those given in Figure 1B, **D**, and **E**, as well as Figure 2B and D and, are shown here for comparison to the values of *mlo* single and double mutants. This Supplemental Figure supports Figures 1 and 2 in the main text.

**Supplemental Figure 2.**
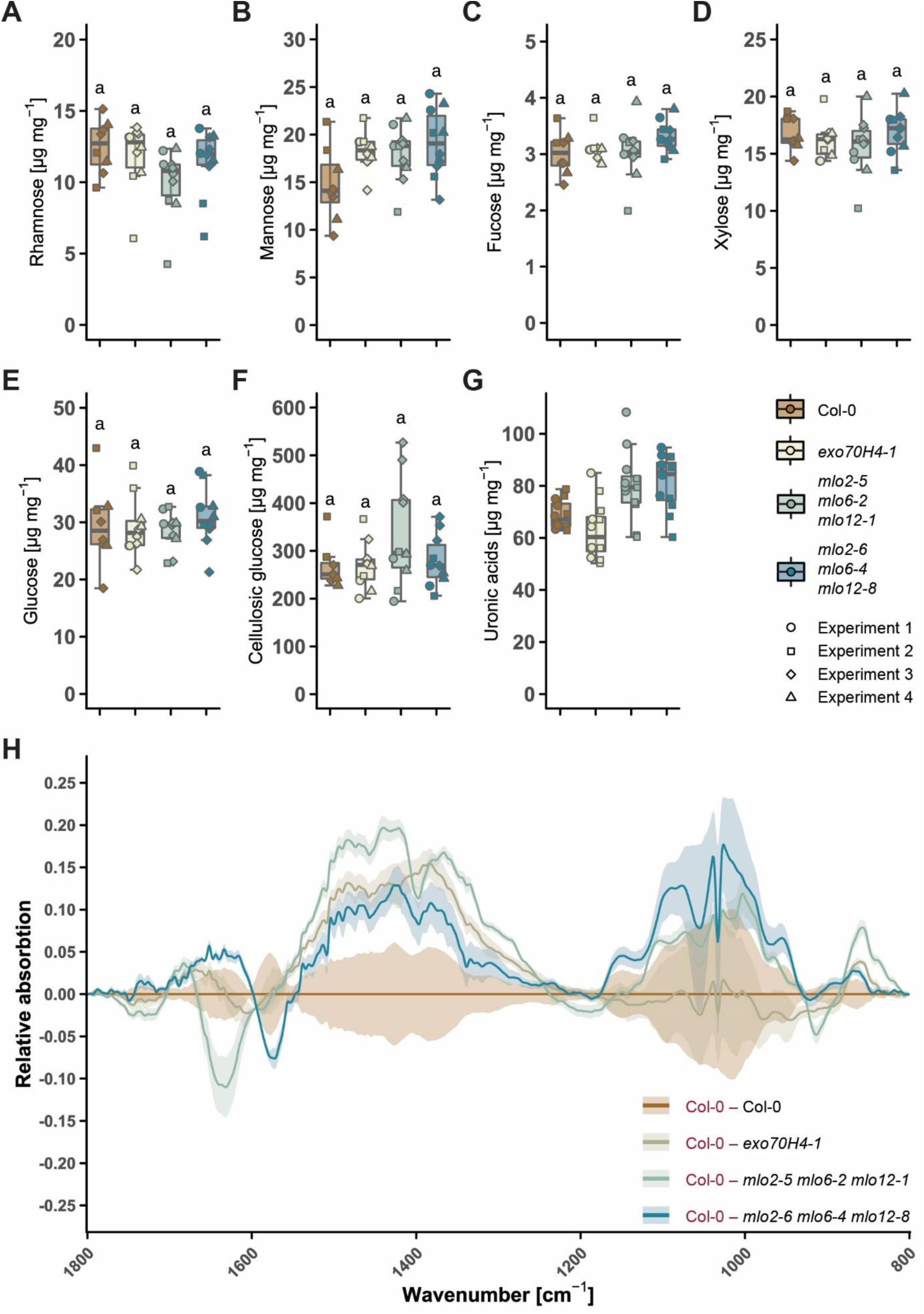
*A. thaliana exo70H4* and *mlo2 mlo6 mlo12* triple mutants have altered trichome cell wall characteristics. **A–G**) Neutral monosaccharide and uronic acid content in trichomes of Col-0 WT and the indicated mutants. The abundance of the neutral monosaccharides rhamnose (**A**), mannose (**B**), fucose (**C**), xylose (**D**), and glucose (**E**) as well as the cellulosic glucose (**F**) content in the alcohol-insoluble residue (in mg) after the hydrolysis of the cell wall matrix of *A. thaliana* rosette leaf trichomes is shown. The boxplots indicate the outcome of at least three independent experiments with two to four replicates each. Letters assign differences of statistical significance (pairwise Student’s *t*-test corrected by FDR, *α* = 0.01). **G**) Uronic acid amount in the trichome dry mass (in mg) of the various genotypes. The boxplot indicates the outcome of two independent experiments with six replicates each. **H**) FTIR difference spectra of various mutant genotypes in relation to Col-0. Each line represents the mean difference spectrum of one experiment with four replicates each. The shaded areas indicate the standard deviation. Minimal turning points signify an increase of the component associated with the respective wavenumber compared to Col-0 whereas peaks imply a reduction. This Supplemental Figure supports Figure 4 in the main text.

**Supplemental Figure 3.**
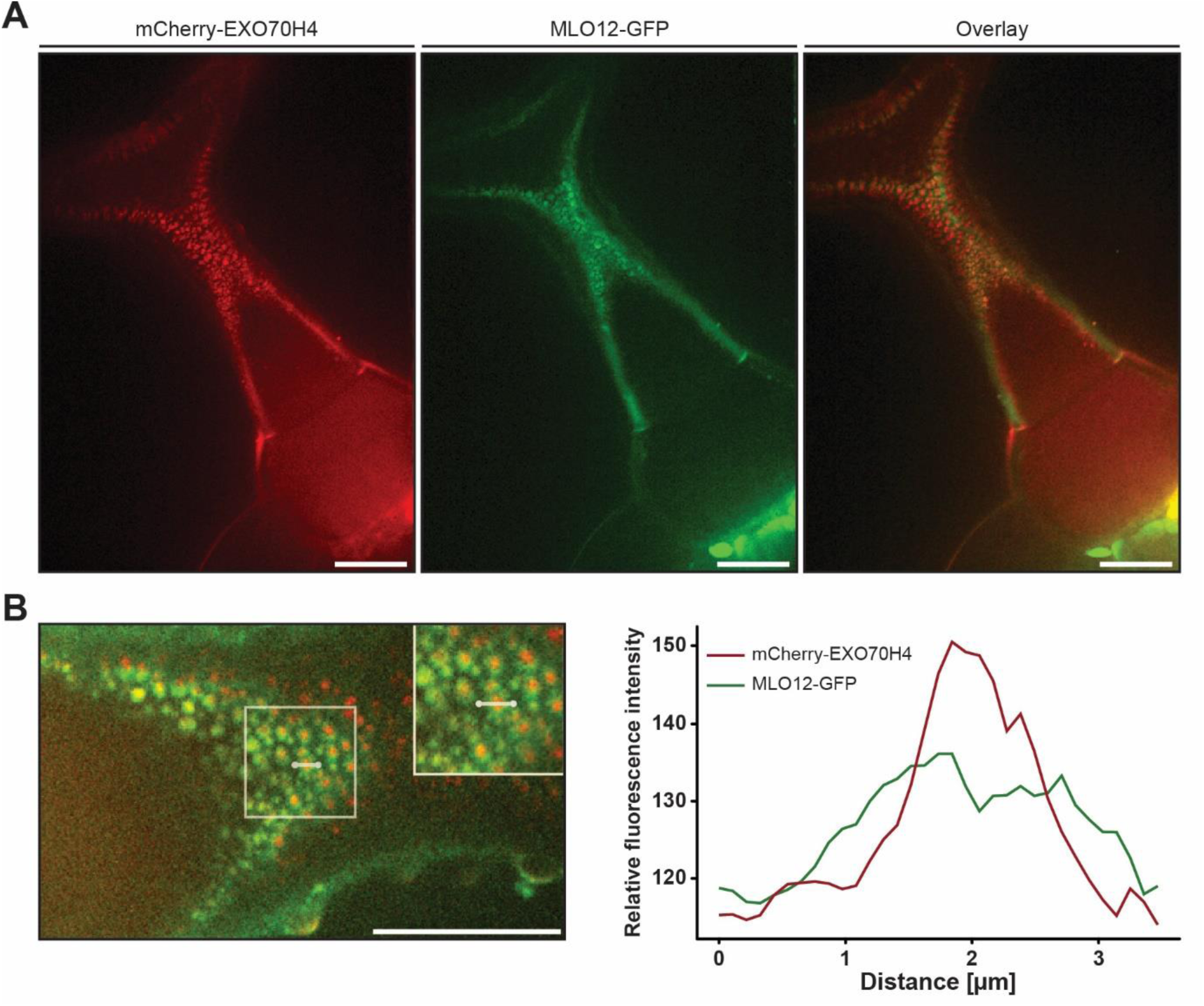
Co-localization details of mCherry-EXO70H4 and MLO12-GFP in trichomes of transgenic lines. **A)** Spinning disc micrographs illustrating the trichome subcellular localization of mCherry-EXO70H4 (left) and MLO12-GFP (middle), expressed in a transgenic *A. thaliana* plant under control of the *EXO70H4* promoter. An overlay image is shown on the right. **B)** Close-up overlay image of a trichome branching area of the transgenic line described in (A) (left panel) and a corresponding fluorescence intensity plot (right panel) for the path marked with a white line in the boxed region (see also enlarged inset in micrograph). Scale bars represent 20 µm. This Supplemental Figure supports Figure 5 in the main text.

**Supplemental Figure 4.**
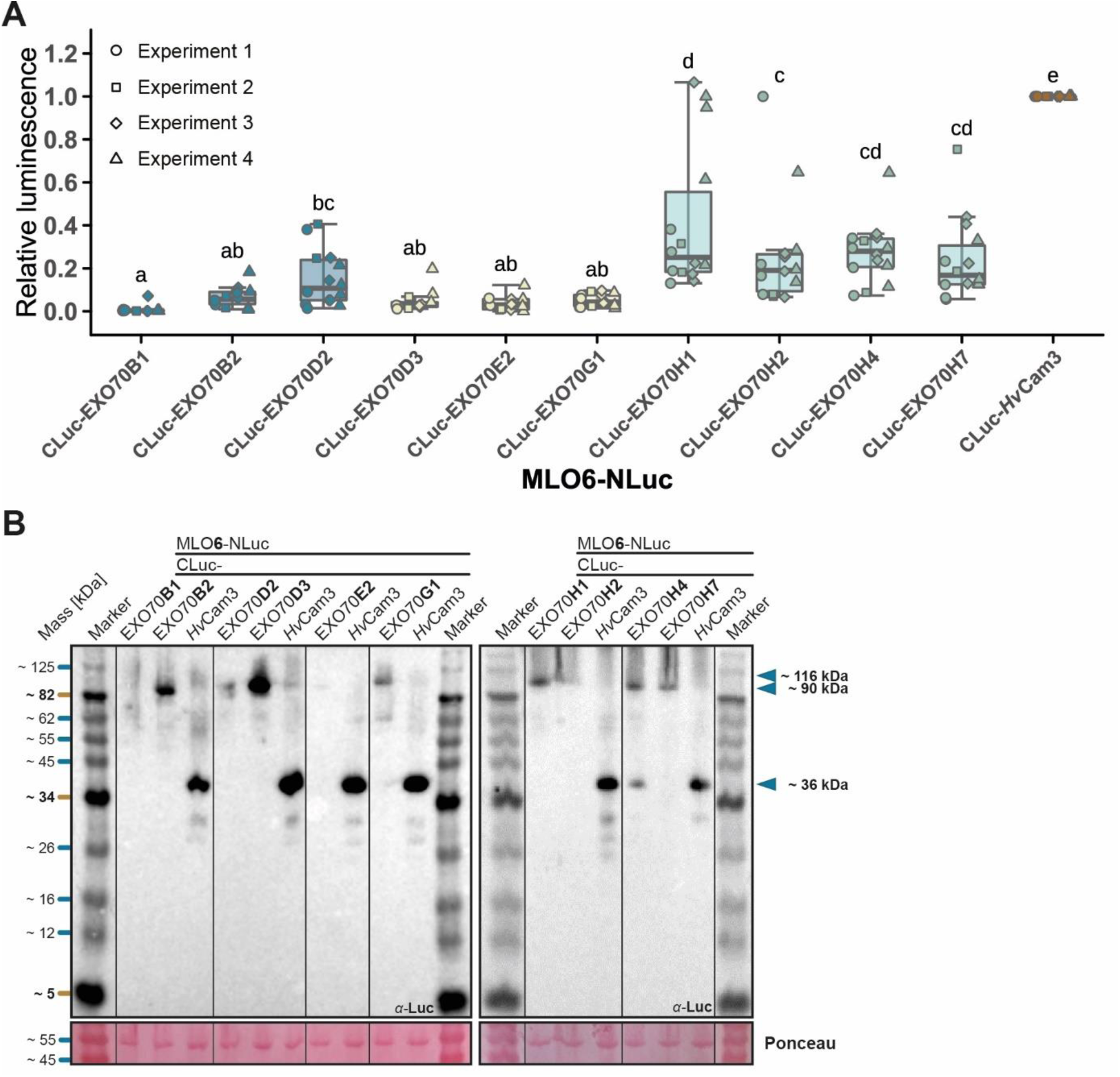
Analysis of the interaction of MLO6 with various EXO70 proteins by LCI. **A**) Luciferase complementation imaging was carried out by transient gene expression in *N. benthamiana* as described in the Materials and Methods section. For normalization, luminescence signals were divided by the signal retrieved for the interaction of MLO6-NLuc with CLuc-*Hv*Cam3 (barley calmodulin 3) on the same leaf (positive control). The boxplot illustrates the outcome of four independent experiments with two to four replicates each. Letters assign differences of statistical significance (pairwise Student’s *t*-test corrected by FDR, *α* = 0.05). **B**) Recombinant luciferase-tagged proteins were detected by immunoblot analysis with an *α*-luciferase antibody. Pooled leaf discs of one independent experiment were subjected to protein extraction. The numbers on the right indicate the predicted molecular mass of the proteins of interest (MLO6-NLuc ∼ 116 kDa, CLuc-EXO70 ∼ 90 kDa, CLuc-*Hv*Cam3 ∼ 36 kDa). Ponceau S staining was carried out to validate equal loading of the gels. This Supplemental Figure supports Figure 6 in the main text.

**Supplemental Figure 5.**
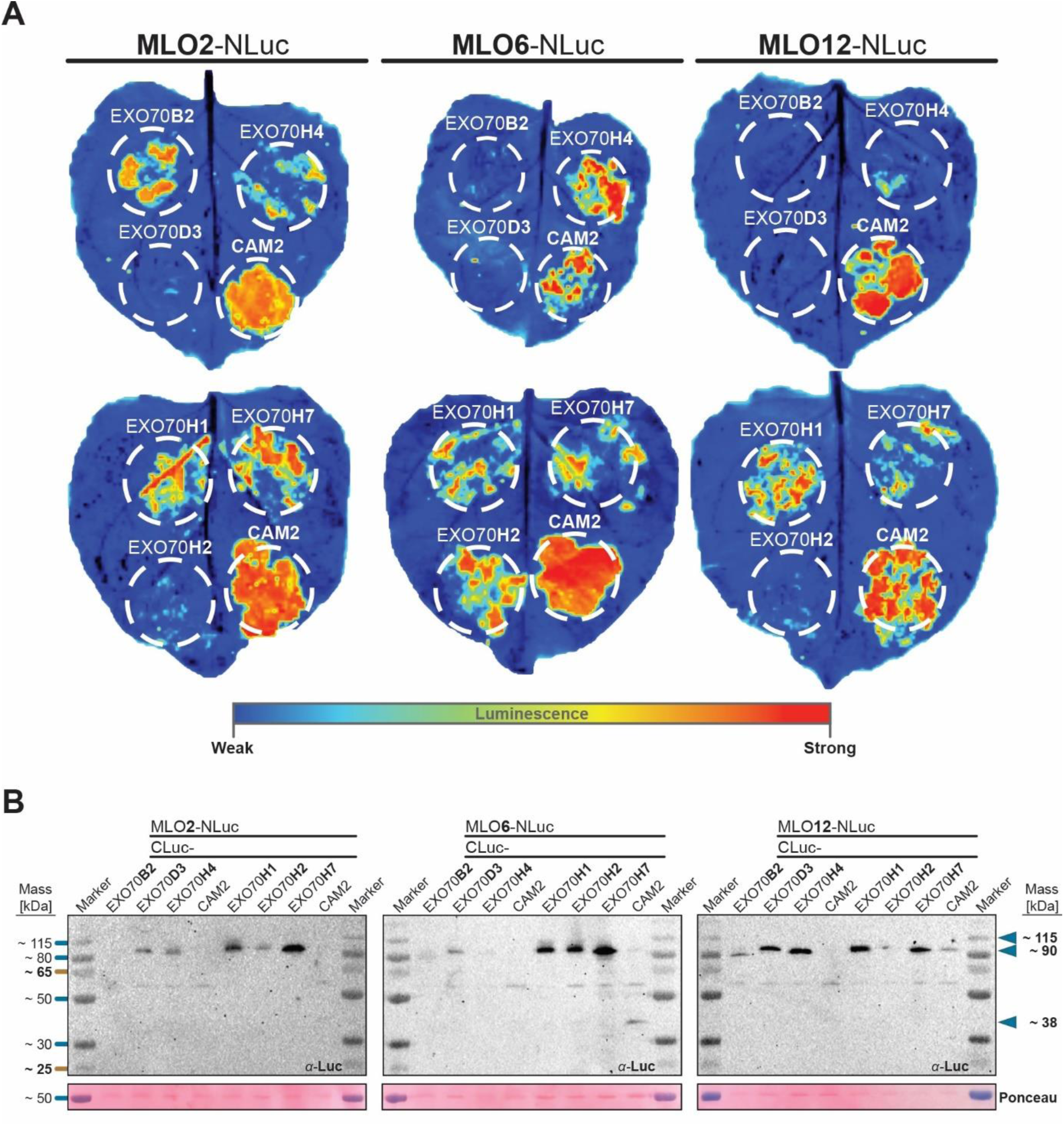
Representative luminescence signals and validation of protein production in leaves of *N. benthamiana* after luciferase complementation imaging. A) Representative pseudo-colored images of *N. benthamiana* leaves after detection of bioluminescence. Luminescence detection was carried out as described in the Materials and Methods section. Luminescence intensity is indicated by the color scale at the bottom of the panels. Leaves are representative of at least four independent experiments with two replicates each, yielding a similar outcome. **B**) Recombinant luciferase-tagged proteins were detected by immunoblot analysis with an *α*-luciferase antibody (upper panels). Pooled leaf discs of one independent experiment were subjected to protein extraction. The predicted molecular mass of the proteins of interest is indicated by the arrowheads and numbers on the right (MLO-NLuc ∼115 kDa, CLuc-EXO70 ∼90 kDa, CLuc-CAM2 ∼38 kDa). Ponceau S staining was carried out to validate equal loading of the gels (lower panels). This Supplemental Figure supports Figure 6 in the main text.

**Supplemental Figure 6.**
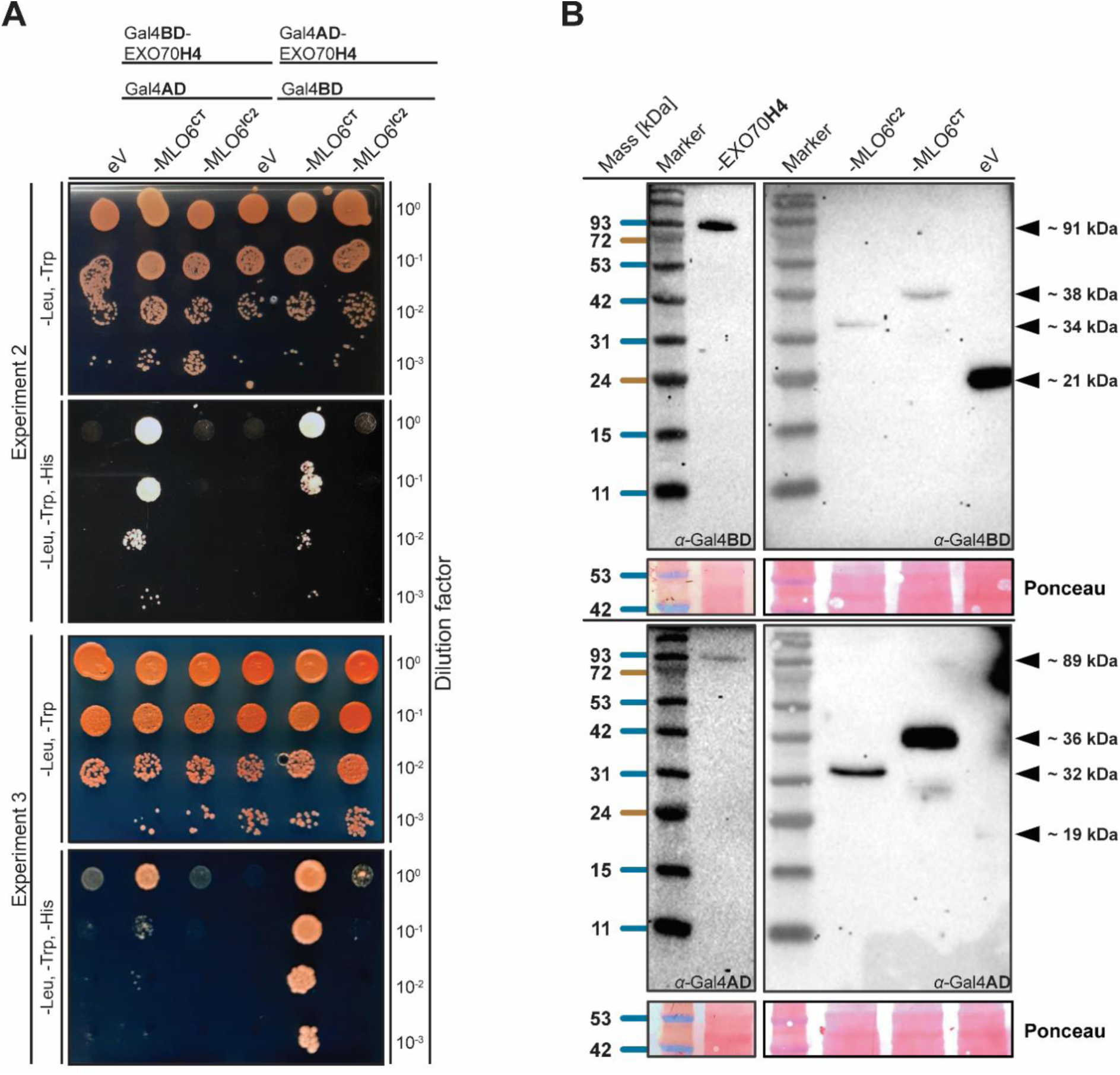
Yeast two-hybrid assays suggest an interaction of EXO70H4 with the carboxyl terminus MLO6. **A**) *S. cerevisiae* cultures expressing recombinant *EXO70H4* and *MLO6* constructs were dropped on a medium without (1) leucine (Leu) and tryptophan (Trp) or (2) without leucine, tryptophan, and histidine (His) in different serial dilutions. An empty vector (eV) encoding the respective tag without MLO6 was used as a negative control. The pictures represent the outcome of three independent experiments, the first of which is shown in Figure 6F. **B**) Immunoblot detection of the binding domain (BD) or the activation domain (AD) of the yeast transcriptional regulator Gal4 that were attached to EXO70H4 or domains of the MLO6 protein, respectively. Molecular masses on the right indicate the predicted mass of the proteins of interest. Ponceau S staining was carried out to validate equal loading of the gels. IC2, second intracellular loop; CT, carboxyl terminus. This Supplemental Figure supports Figure 6 in the main text.

